# Human cytomegalovirus regulates host DNA repair machinery for viral genome integrity

**DOI:** 10.1101/2025.10.18.683233

**Authors:** Pierce Longmire, Sebastian Zeltzer, Kristen Zarrella, Olivia Daigle, Marek Svoboda, Justin M. Reitsma, Scott S. Terhune, Carly Bobak, Giovanni Bosco, Felicia Goodrum

## Abstract

The DNA damage response (DDR) encompasses a multitude of interconnected pathways that serve as a cellular defense to protect genome integrity. Dysregulation or failure of these pathways results in cancers and genetic disease. DNA viruses, including the herpesvirus cytomegalovirus (CMV), activate DDR signaling during their replicative program. The mechanisms by which they commandeer these responses for replication of their genome is less well understood. Here, we define a viral protein, UL138, that modulates the activity of host DDR pathways. The loss of UL138 results in structural variants, including inversion, deletions, duplication, with signature of homology-directed repair and other DDR pathways. The actions of UL138 are due, in part, to its modulation of pathways regulated by the cellular deubiquitinating complex that targets proliferating cell nuclear antigen (PCNA) and Fanconi Anemia effectors, FANCD2 and FANCI. However, we also show that UL138 accesses pathways independent of USP1-PCNA/FANCD2/FANCI. Disruption of UL138 or these pathways impacts viral genome replication and had consequences for viral genome integrity. This work provides mechanistic insight into the long-standing questions of how DNA viruses recruit, modulate and use cellular DDR pathways. It also puts forth CMV as a model system for further defining these pathways in human cells.

**GRAPHICAL ABSTRACT:** 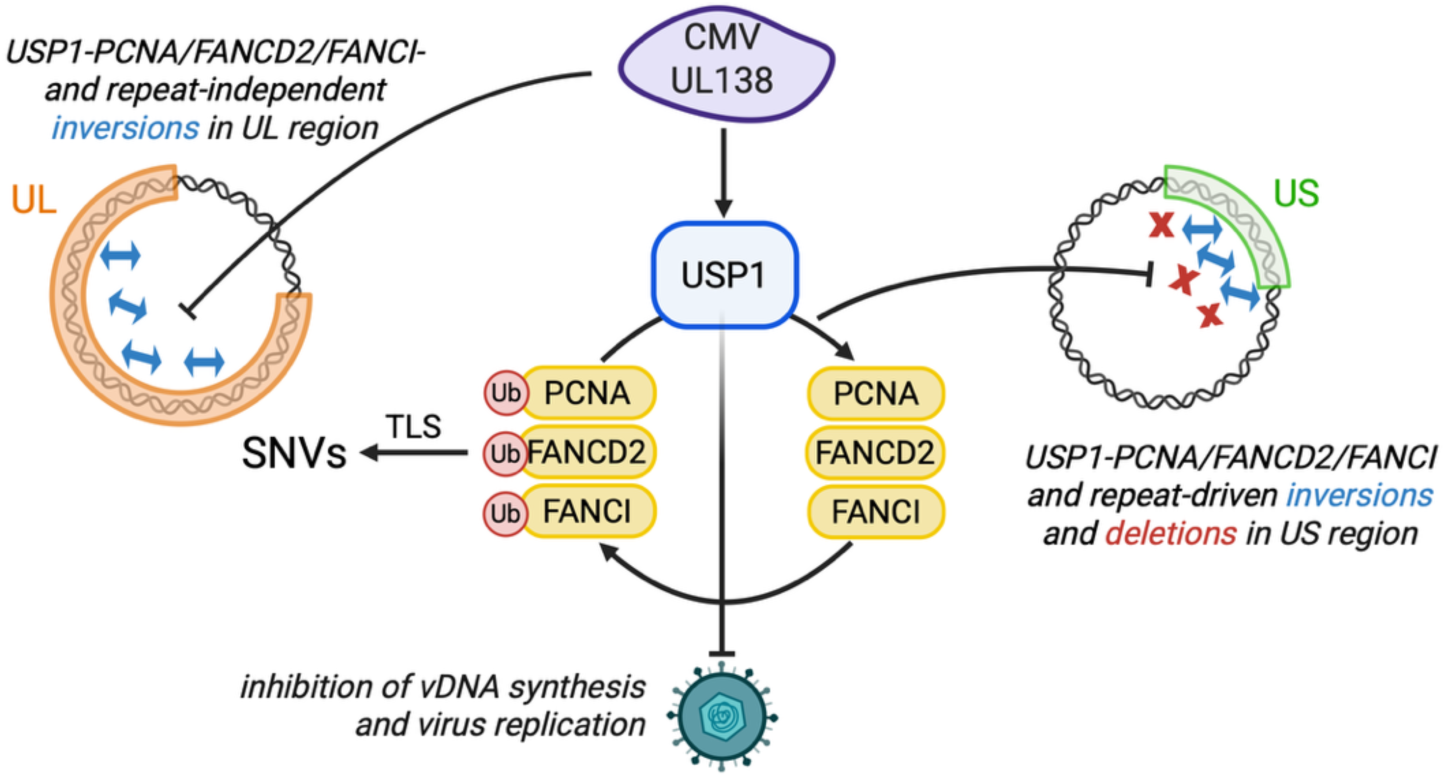

**Major findings:**

- UL138 protects HCMV genome integrity through its interaction with USP1-PCNA/FANCD2/FANCI and repeat sequence-mediated pathways
- UL138 protects HCMV genome integrity in the UL region by mechanisms independent of repeat sequences and USP1-PCNA/FANCD2/FANCI
- Collectively, US and UL regions of the HCMV genome have fundamentally different pathways acting on them and UL138 modulates DNA damage and repair pathway choice.

## INTRODUCTION

The DNA damage response (DDR) and subsequent repair pathways represent an essential cellular defense against mutations that can compromise genome integrity and lead to a variety of human diseases, including cancers. Repair pathways are also an important defense against foreign genetic material, such as that of viruses, which pose a threat to host genomic stability (1). While DNA viruses have evolved tactics to suppress many aspects of this defense system, they commandeer other features to replicate their own genomes (2–4). Host DDR factors may also be utilized in the establishment of persistent infections to circularize and maintain viral genomes for latency (5). However, we lack a full understanding of how these virus-host interactions contribute to virus replication, especially for viruses with large, complex DNA genomes.

Human cytomegalovirus (CMV) is a pervasive betaherpesvirus that latently infects a majority of the global population. With a large, double-stranded DNA genome of over 236-kilobases that is estimated to encode more than 200 gene products, CMV is the largest known human herpesvirus (6–8). The CMV genome is complex, with high G-C content that support G-quadruplex structures and repetitive sequences, all of which provide challenges for its replication (9). CMV induces a DNA damage response upon infection and recruits several host DNA repair factors to nuclear viral replication compartments (RCs), sites of viral DNA (vDNA) synthesis and transcription, during productive infection (10–12). While these observations may simply represent a host response to infection, they may be inhibited or redirected by the virus for the viral life cycle. The requirement for ATM in virus replication suggests the importance of homology-directed repair (HDR) pathways for the replication of the viral genome (10). However, the specific pathways targeted, the mechanisms by which they are altered or recruited, and the viral factors involved remain largely uncharacterized. Even less is known about the role of host DNA repair pathways in the regulation of viral latency and reactivation, which poses serious disease and mortality risk for immunocompromised individuals. Complex interplay between viral determinants and host biology is central to the regulation of viral gene expression, genome amplification, and the decision to establish latency or replicate.

We previously showed that CMV recruits specialized host DNA polymerases from the translesion synthesis (TLS) pathway during productive infection to modulate virus replication and maintain viral genome integrity (13). TLS is a DNA damage bypass pathway that may occur when the replication complex encounters DNA lesions, such as UV-induced pyrimidine dimers, that stall or block DNA fork progression (14). TLS polymerases are low-fidelity and, while they minimize the risk of collapse of stalled forks, they are more likely to introduce mutations during this step compared to replicative DNA polymerases (15). TLS polymerases also have other more poorly defined roles in HDR (16–18). The mechanisms by which TLS polymerases are recruited and regulated in CMV infection remain undefined. Loss of TLS polymerases in infection decreased single nucleotide variants (SNVs), indicative of canonical TLS function, but also increased structural variants, particularly inversions, which are indicative of non-canonical TLS functions such as homology-directed repair pathways. TLS polymerases canonically work in concert with the DNA clamp and processivity factor, proliferating cell nuclear antigen (PCNA), in repair of cellular DNA. However, we have shown that TLS polymerases also have PCNA-independent roles in the context of CMV infection (19). In response to damage, PCNA is monoubiquitinated (mUb) to regulate DNA repair, most notably repair involving TLS polymerases (20,21). Deubiquitination of PCNA downregulates TLS polymerase recruitment and is mediated by the deubiquitinating enzyme ubiquitin specific peptidase 1 (USP1) with its scaffold USP1-associated factor 1 (UAF1), also known as WD-repeat containing protein 48 (WDR48) (22–24).

The UAF1-USP1 complex is also important for its role in modulating steps in the Fanconi Anemia (FA) repair pathway (23). FA is a hereditary genetic disorder marked by hypersensitivity to DNA crosslinking agents, genomic instability, and increased susceptibility to cancer (25). FA repair includes multiple genes that respond to DNA interstrand crosslinks during replication, including a core E3 ligase complex that monoubiquitinates FANCD2 and FANCI at the site of damage(26). After this critical step, the mUb-FANCD2/FANCI complex recruits nucleases, polymerases, and other enzymes to repair the crosslinked DNA, converging with various other repair pathways to repair the lesion (27). Beyond DNA crosslink repair, the FA pathway is implicated in the response to replication stress by protecting stalled forks from degradation (26). Through its deubiquitinating activity, UAF1-USP1 plays an important role in termination of this repair pathway.

We recently identified an interaction between UL138, a determinant of CMV latency, and the UAF1-USP1 complex and showed that it functioned to sustain activation of signal transducer and activator of transcription 1 (STAT1) signaling (28). This interaction is important to viral latency as either chemical inhibition of USP1 or disruption of UL138 permits CMV to replicate in CD34+ hematopoietic progenitor cells, thus preventing the establishment of latency (28). Given the additional role of UAF1-USP1 regulating host DDR pathways, we hypothesized that UL138 might commandeer the host DDR through its interaction with UAF1-USP1 and impact stability of the viral genome. Here we show that USP1, PCNA, and FANCD2, were re-localized to sites of viral DNA synthesis during productive CMV infection in primary fibroblasts. While UAF1 facilitates virus replication, USP1 and its substrates restrict CMV replication, suggesting a role for other UAF1-USP complexes in virus replication. UL138-UAF1-USP1 stimulates deubiquitination of PCNA and FANCD2 during infection with restrictive effects on viral genome synthesis. Strikingly, disruption of UL138 or depletion of USP1 and its substrates induced viral genome rearrangements, highlighting an intriguing and novel function for UL138 and the UAF1-USP1 interaction in CMV replication and genome integrity. While increased SNVs is a signature of canonical TLS bypass functions, structural variants, particularly those dependent on repetitive sequences, are a signature of non-canonical TLS function and are suggestive of HDR mechanisms. Intriguingly, some rearrangements, particularly in the UL region of the genome, occur independently of USP1 and its substrates and are likely independent of TLS polymerases.

Altogether, we provide mechanistic insights into how CMV directly commandeers host DNA repair pathways to control CMV replication. Moreover, CMV provides a novel model for investigating DDR pathways that remain poorly understood in human cells.

## MATERIALS AND METHODS

### Cells and Viruses

Primary human lung MRC-5 fibroblasts (ATCC CCL-171) were maintained in DMEM containing 10% fetal bovine serum, 10 mM HEPES, 1 mM sodium pyruvate, 2 mM L-alanyl-glutamine, 0.1 mM nonessential amino acids, 100 U/mL penicillin, and 100 µg/mL streptomycin. Cells were infected with the low-passage CMV strain, TB40/E, a gift from Dr. Christian Sinzger, which was engineered to expressed green fluorescent protein from the SV40 promoter. TB40/E-Δ*UL138*_STOP_ was previously characterized (29).

### RNAi

All siRNAs used in this study were purchased as a SMARTpool from the ON-TARGETplus siRNA line by Dharmacon (Horizon Discovery): non-targeting control (NTC), UAF1, USP1, FANCD2, FANCI, and PCNA. To achieve knockdown, siRNAs were reverse-transfected into cells using RNAiMAX transfection reagent (Thermo Fisher Scientific) as described in (30). Briefly, 100 pmol of siRNA was mixed with 750 µL Opti-MEM. In a separate tube, 30 µL of RNAiMAX (6 volumes relative to siRNA) was mixed with 750 µL Opti-MEM. These two mixtures were thoroughly combined, spread evenly on a sterile 10 cm dish, and incubated at room temperature for 5 minutes. Finally, approximately 2 million cells were then added to the dish in 10 mL growth media (10 nM siRNA final). Cells were incubated at 37°C and after two days of growth-arrest, cells were infected with CMV as described.

### RT-qPCR and qPCR

Cells were lysed in DNA/RNA lysis buffer and DNA and RNA were isolated from cells using Quick-DNA/RNA Miniprep Kit (Zymo Research) according to the manufacturer’s protocol. For RT-qPCR, cDNA was synthesized from RNA using Transcriptor First Strand cDNA Synthesis Kit (Roche) and quantified using the ΔΔCt method relative to the cellular housekeeping control, H6PD. Viral genomes were quantified from DNA by qPCR using PowerUp SYBR Green Master Mix (Applied Biosystems) and primers specific to the HCMV genomic region encoding the β2.7 transcript. Absolute genome copy number was calculated using a TB40/E BAC standard curve and normalization to cellular RNaseP. Reactions were run on a QuantStudio 3 system (Applied Biosystems; ThermoFisher Scientific) using QuantStudio Design & Analysis Software (v1.4.3).

### Immunofluorescence

Fibroblasts were seeded onto 12mm glass coverslips (Electron Microscopy Sciences) in 24-well plates. The following day cells were mock infected or infected with TB40/E-WT virus at an MOI of 1. At 24, 48, and 72 hpi, cells were washed in cold cytoskeletal buffer for 2 minutes on ice as previously described. Cells were then washed with PBS and fixed in 2% paraformaldehyde for 20 minutes at room temperature (for USP1 and FANCD2 staining) or in 100% methanol for 10 minutes at -20°C (for PCNA staining). Coverslips were then processed for indirect immunofluorescence as previously described (13). Briefly, coverslips were in blocking buffer containing normal goat serum, bovine serum albumin, and human serum for one hour at room temperature. Proteins of interest were detected using specific primary antibodies for one hour at room temperature or overnight at 4°C as described in Table S1. Coverslips were washed 3 times in PBS + 0.05% Tween20 and then incubated in secondary antibodies (AlexaFluor 546, AlexaFluor 647, or AlexaFluor 488 goat anti-mouse or goat anti-rabbit [Invitrogen]) for 30 minutes at room temperature. Coverslips were incubated in DAPI for 5 minutes and then washed three times in PBS + 0.05% Tween20. Finally, coverslips were mounted onto microscope slides (Fisher Scientific) using Prolong Gold Antifade Mounting Reagent (Invitrogen).

### Deconvolution microscopy

Images were obtained using a DeltaVision RT Inverted Deconvolution Microscope, 60x objective. Representative single plane images with 0.2 µM thickness were adjusted for brightness and contrast using Fiji/ImageJ software.

### Immunoblotting

Whole cell lysates were extracted using RIPA lysis buffer (Pierce) and manual scraping. 50 µg of lysate was loaded onto precast 4-12% bis-tris gels (ExpressPlus™; GenScript) and then proteins were separated by electrophoresis and transferred onto 0.45 µm pore size PVDF membranes (Immobilon®-FL; Millipore). Proteins of interest were detected using primary antibodies described in Table 1 and fluorophore-conjugated secondary antibodies. Images were obtained using a Li-Cor Odyssey CLx scanner and protein levels were quantitated using Image Studio Lite software.

**Genomic sequencing and computational analysis**

Fibroblasts were seeded onto 6-cm dishes and reverse-transfected with a pool of siRNAs as described above. Five technical replicates were seeded for each condition. At two days post-transfection, cells were infected with CMV TB40/E-WT or -Δ*UL138*_STOP_ at an MOI of 1. Virus inoculum was removed at 2 hpi and cells were provided fresh media. At 96 hpi, when maximal cytopathic effect (CPE) was observed, cells were washed with PBS and collected in DNA lysis buffer containing 200 µg/mL proteinase K by manual scraping. After a two hour 55°C incubation for proteinase K digestion, cellular and viral DNA were isolated using phenol-chloroform extraction. DNA was similarly extracted from the virus stock used for infection. All purified DNA was submitted to SeqCenter (Pittsburgh, PA).

Each sample (including virus stocks) was sequenced on a NextSeq 2000 Illumina short read sequencer to yield paired-end short reads. The sequencing reads were aligned to the human reference genome GRCh38 (31) using Bowtie2 v2.5.1 (32) with the following parameters: --very-sensitive for sensitive alignment, --seed 1 for seeding alignment. After alignment, reads aligned to the human genome were filtered out using Samtools v1.17 (33). The following parameters were used: -f UNMAP,MUNMAP to extract reads that were not aligned to the human genome, and -bh to output the filtered alignments in BAM format. CMV junctions and SNVs were then detected using a two-pass analysis to ensure comparable sequencing coverage between the samples. Non-human reads were first aligned to the reference CMV genome with breseq v0.38.1(34) using polymorphism-prediction mode. Using the mean sequencing coverage of reads aligned to the reference from the first breseq run output, the sequencing reads from each sample were subsampled with seqtk v1.4q (https://github.com/lh3/seqtk) using parameter -s 100 and the appropriate respective proportions to yield the same mean coverage equal to that of the sample with the lowest coverage (WT siNTC). The subsampled FASTQ files were then used to detect the junctions and SNVs by running breseq in the --polymorphism-prediction mode again. Subsequent data analyses were conducted in R version 4.3.2 (35). The novel junctions and SNVs were obtained by removing any of those found in the TB40/E-WT virus stock sample from the total junctions and SNVs detected in each experimental sample. A Poisson two-sided test was used to compare junction frequency across experimental conditions, with a junction frequency cutoff of 0.025 employed to refine the selection of relevant sequences. Additionally, circle plots were constructed using ‘circlize’ (version 0.4.16) (36) to visualize the locations and relationships of these sequences, providing a comprehensive view of their distribution and interaction within the genome. This multifaceted approach allowed for a nuanced exploration of genomic junctions, highlighting significant variations and patterns across experimental conditions.

### Statistical analysis

All statistics were analyzed using GraphPad Prism version 9 software. Specific statistical tests are indicated in figure legends and included paired and unpaired Student’s t-tests, one-way analysis of variance (ANOVA), and two-way ANOVA with multiple comparisons tests where applicable. Asterisks represent statistically significant differences based on P-values from these tests: * P <0.05, ** P <0.01, *** P <0.001, ****P <0.0001.

## RESULTS

### UAF1 facilitates and USP1 suppresses CMV replication

Our previous studies demonstrated a requirement for USP1 activity in the establishment of latent CMV infection (28). Disruption of USP1 activity with the chemical inhibitor C527 results in a failure of CMV to establish latency in CD34+ HPCs, suggesting that, like UL138, USP1 restricts CMV replication (28). Importantly, inhibition of USP1 with C527 in UL138-mutant virus infection does not further impact virus replication. To further explore the role of USP1 in virus replication, we depleted USP1 or its scaffold, UAF1, in primary fibroblasts to assess productive CMV replication. Cells were growth-arrested by contact inhibition such that depletion of host factors important for cellular DNA synthesis and repair does not induce stress. Growth-arrested cells readily infect and support the HCMV replicative cycle. To understand specific contributions of UL138, cells in which UAF1 or USP1 had been knocked down were then infected with TB40/E-GFP virus (wild-type, WT) or a recombinant virus in which UL138 protein expression is disrupted through insertion of a premature stop codon on the 5’ end of *UL138* (Δ*UL138*_STOP_) (29). siRNA knockdown of USP1 increased viral genome copy number ∼2-fold (Fig. 1A) and virus progeny yields ∼5-fold (Fig. 1B) compared to non-targeting control (NTC) siRNAs in WT-infected fibroblasts. This finding is consistent with our previous findings that USP1 restricts vDNA synthesis and subsequent production of progeny virus (28). Of note, USP1 knockdown increased vDNA synthesis and virus yield in Δ*UL138*_STOP_ infection, but the increase in yield was not statistically significant. Confirmation of knockdown is shown in Figure 1C.

**Figure 1.**
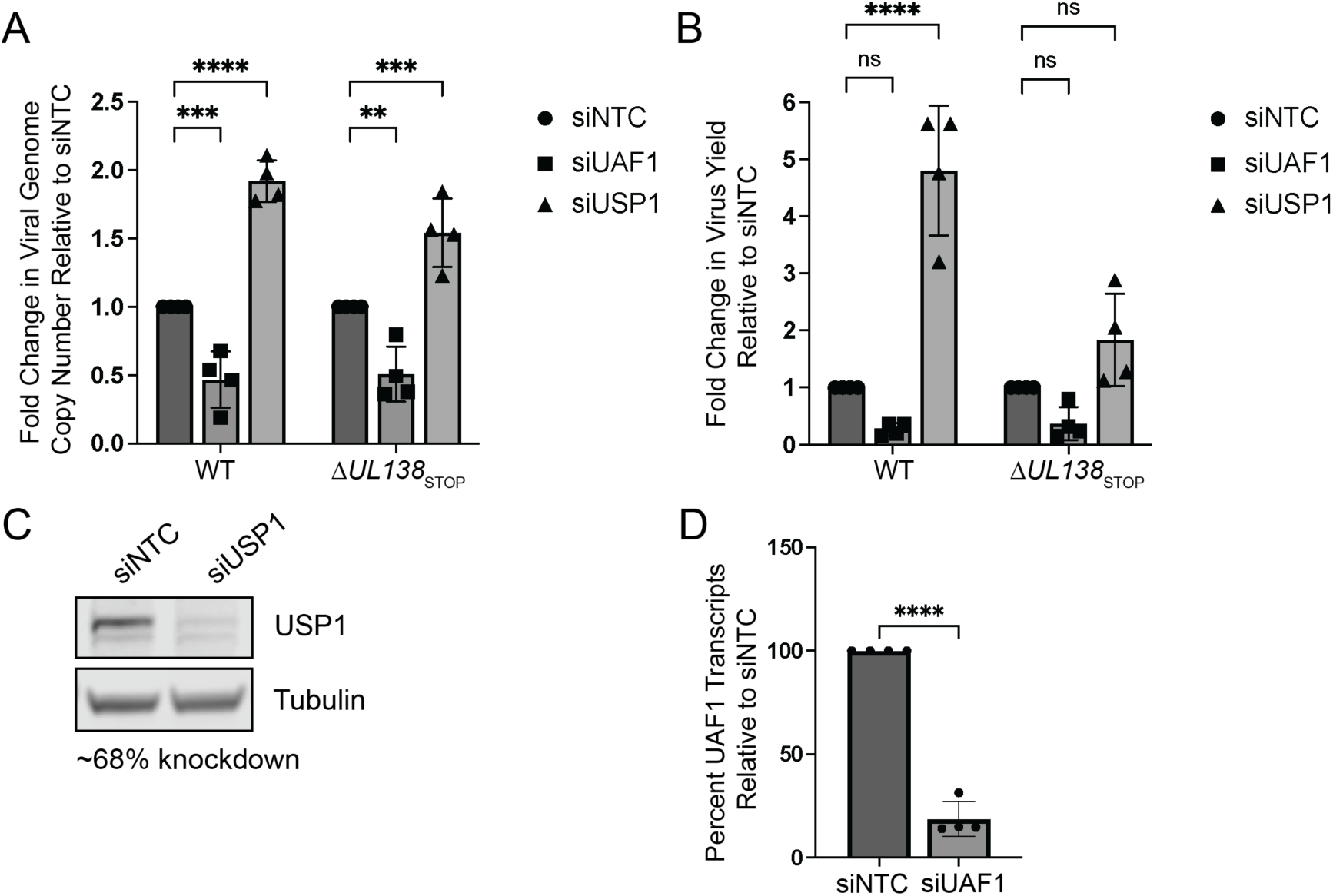
USP1 restricts CMV replication. MRC-5 fibroblasts were reverse transfected with a siRNA pool targeting UAF1 or USP1 or a non-targeting control (NTC). At two days post-transfection, cells were infected with CMV TB40/E-GFP wild-type (WT) or TB40/E-Δ*UL138*_STOP_ at an MOI of 1 and cells were harvested at 96 hpi. (A) Viral genome copy number was determined by qPCR using a TB40/E BAC standard curve and primer set designed for the region of the viral genome encoding the β2.7 transcript. The graph represents the fold change in viral genome copy number relative to siNTC. (B) Virus yields were measured by TCID_50_ and normalized relative to the WT siNTC condition. For A-B statistical significance was determined by two-way ANOVA with Dunnett’s multiple comparisons test. Asterisks (**P <0.01, ***P <0.001, ****P <0.0001) represent statistically significant differences determined for four independent experiments. ns, non-significant. (C) Whole cell lysates were collected at two days post-transfection to determine efficiency of USP1 knockdown by immunoblotting using monoclonal antibodies for USP1 and tubulin as a loading control and secondary antibodies conjugated to DyLight™ 680 (mouse) or 800 (rabbit). Densitometry analysis of three biological replicates reveals an average 68% knockdown compared to siNTC. (D) At two days post transfection, RNA was collected for RT-qPCR to determine the efficiency of UAF1 knockdown relative to siNTC. H6PD was used as a housekeeping control for cell number. Over all experiments, an average knockdown efficiency of 81% was achieved. A paired *t* test was used to determine statistical significance, ****P <0.0001, over multiple independent experiments.

Enzymatic activity of USP1 is strongly stimulated through interaction with its scaffold and cofactor, UAF1 (23). UAF1 also serves as a scaffold for other deubiquitinating enzymes, USP12 and USP46 (28,37). siRNA depletion of UAF1 reduced genome synthesis in both WT and Δ*UL138*_STOP_ infection (Fig. 1A), consistent with one other study (38). However, there was not a corresponding reduction in viral titers (Fig. 1B). UAF1 RNA levels were analyzed to confirm knockdown (Fig. 1D). As UAF1 is also important for activities of USP12 and USP46, this finding suggests differential roles for UAF1-USP complexes in CMV infection beyond activating USP1 (28,37).

### CMV re-localizes USP1 to replication compartments

Consistent with its role in DNA repair, USP1 localizes to the nucleus (39,40). CMV has been shown to recruit various host factors, especially many involved in host DNA synthesis and repair, to viral RCs in the nucleus, sites of viral DNA synthesis and transcription (10,12,13,41). USP1 predominantly localized within the nuclear viral RCs, marked by the viral DNA polymerase processivity factor, pUL44 (Fig. 2) (42,43). This was observed from early (24 hours post infection, hpi) to late (72 hpi) times in infection, supporting a role for USP1 on viral DNA. This localization was not dependent on UL138, as USP1 was also localized to RCs in Δ*UL138*_STOP_ infection (Fig. S1A). At late stages of infection (72 hpi), USP1 was observed in juxtanuclear structures resembling viral assembly compartments (44). These results suggest a role for USP1 in regulating vDNA synthesis. The distinct localization of UL138 (45) and USP1 at early times suggest that USP1 activity may be modulated by UL138 in secretory membranes within the viral assembly compartment, the predominant site of UL138 localization (29,45).

**Figure 2.**
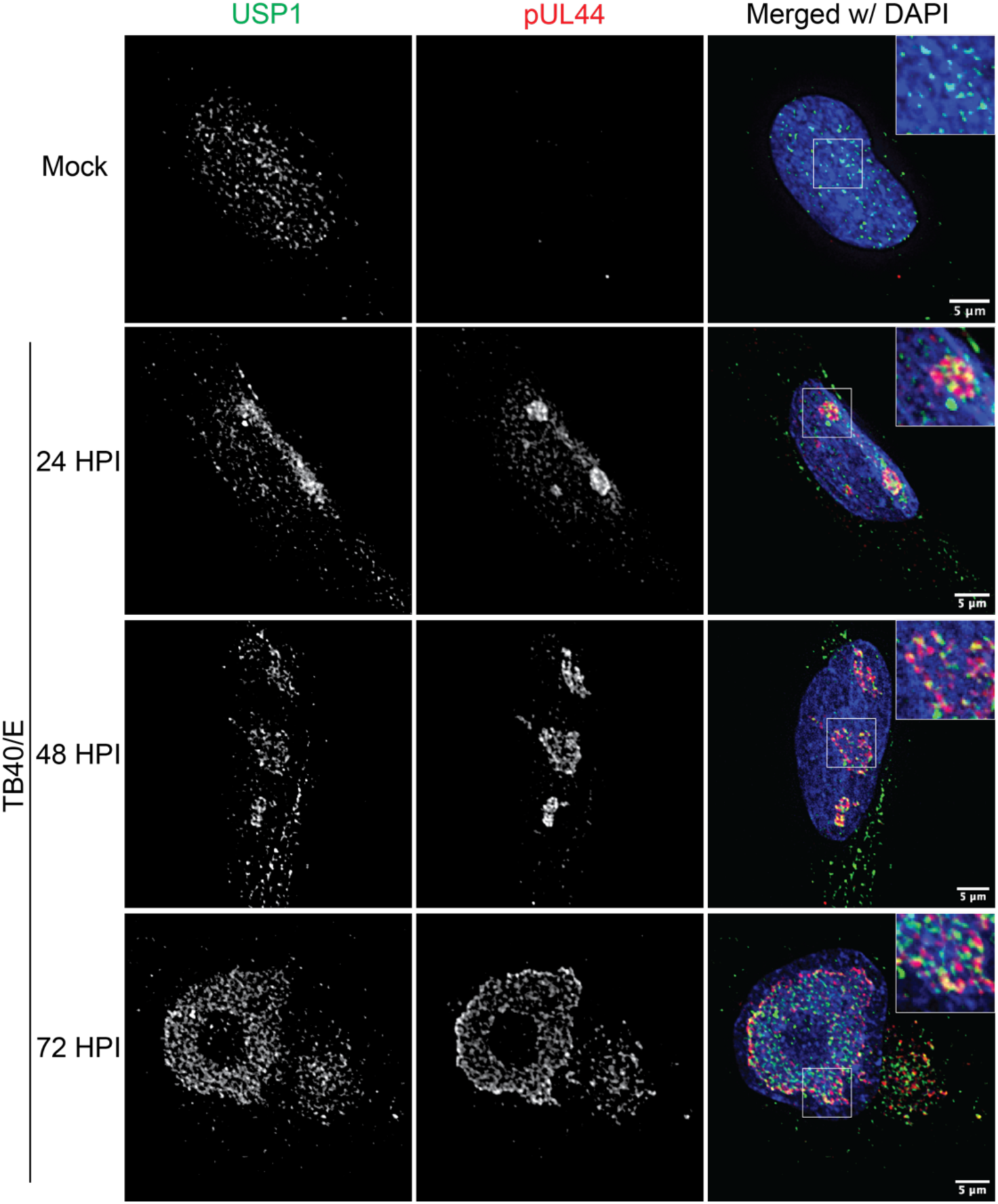
CMV re-localizes USP1 to replication compartments. Fibroblasts were mock-infected or infected with TB40/E-GFP at an MOI of 1. At the indicated time points (24 hpi for mock), cells were fixed and processed for indirect immunofluorescence using monoclonal antibodies to USP1 and pUL44 to mark sites of viral DNA replication and secondary antibodies conjugated to Alexa Fluor® 546 (green) or 647 (red). A magnified image is shown in the top right corner inset of the merged image. DAPI is used to mark the cell nucleus. Images were captured using a DeltaVision deconvolution microscope, and each deconvolved image corresponds to a single focal plane. Scale bar, 5 µm.

### CMV re-localizes PCNA and FANCD2 to replication compartments

We also characterized localization of USP1-regulated DDR proteins in productive infection. Our group and others have previously shown PCNA present in CMV RCs irrespective of virus strain (13,19,46–49). Here we demonstrate that PCNA was re-localized to RCs between 24 and 72 hpi in CMV TB40/E infection (Fig. 3A). FANCD2 also localizes to the nucleus and forms foci at sites of DNA damage (50), and throughout 72 hpi, we found FANCD2 present within RCs marked by pUL44 (Fig. 3B). Because FANCD2 and FANCI interact to nucleate DNA repair (27), we presume that FANCI is present within RCs as well but lack antibodies sufficient for detection. PCNA and FANCD2 were also re-localized to RCs in Δ*UL138*_STOP_ infection (Fig. S1B-C), which suggests this phenomenon occurs independently of UL138, as was observed for USP1. Intriguingly, while USP1, PCNA, and FANCD2 all localize to RCs marked by pUL44, these host proteins only partially overlap with pUL44 localization, indicating that while both viral and host DNA replication proteins are recruited to RCs, they may also function independently. Based on these findings, we propose that these DDR proteins could function in the regulation of vDNA synthesis.

**Figure 3.**
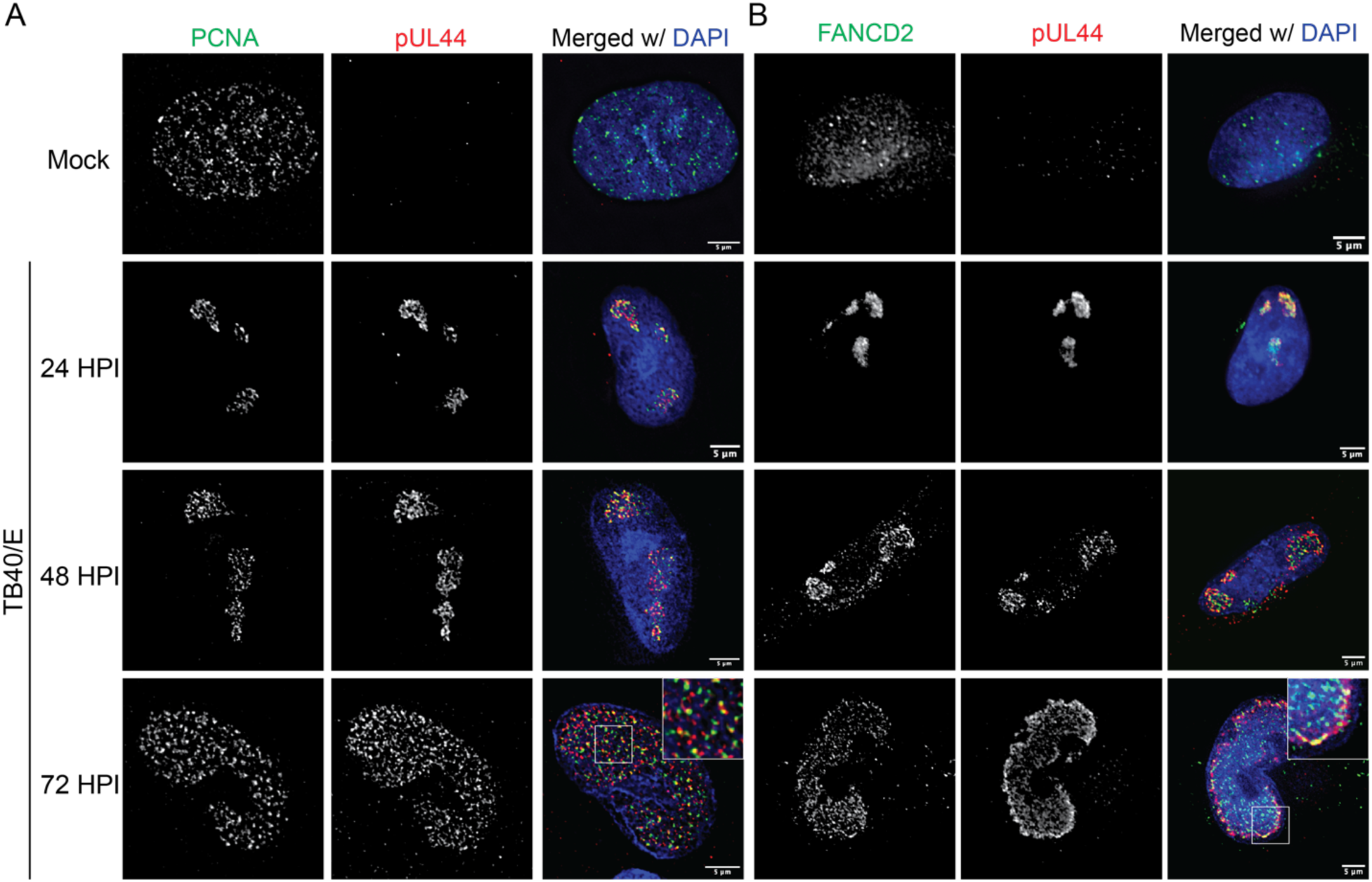
CMV re-localizes PCNA and FANCD2 to replication compartments. Fibroblasts were mock-infected or infected with TB40/E-GFP at an MOI of 1. At the indicated time points (24 hpi for mock), cells were subject to CSK extraction and fixation and then processed for indirect immunofluorescence. (A) PCNA and (B) FANCD2 were detected using monoclonal antibodies specific to each and α-pUL44 to mark sites of viral DNA synthesis and secondary antibodies conjugated to Alexa Fluor® 546 (green) or 647 (red). A magnified image is shown in the top right corner of the merged image at 72 hpi. Images were captured using a DeltaVision deconvolution microscope, and each deconvolved image corresponds to a single focal plane. Scale bar, 5 µm.

### UL138 and USP1 stimulate deubiquitination of PCNA and FANCD2 in CMV infection

To determine if UL138 impacts ubiquitination of UAF1-USP1 substrates, we assessed mUb levels of PCNA and FANCD2 throughout 72 hpi for WT and Δ*UL138*_STOP_ viruses. We previously reported that vDNA synthesis induced mUb-PCNA during CMV infection (13,19). Here, we observed that in the absence of UL138 protein, mUb-PCNA accumulated more rapidly and to greater levels than in WT infection (Fig. 4A, quantified in 4C). In contrast, non-ubiquitinated PCNA levels were maintained throughout either WT or Δ*UL138*_STOP_ infection. To address whether this was specific to the PCNA substrate of USP1, we also assessed protein levels of FANCD2. Similar to mUb-PCNA, we found that mUb-FANCD2 accumulated more rapidly in the absence of UL138 (Fig. 4B, quantified in 4D). Consistent with this, we observed a decrease in the unmodified form of FANCD2. These data suggest a specific role for UL138 in promoting deubiquitination of mUb-PCNA and mUb-FANCD2, likely through interaction with UAF1-USP1. Because mUb of the FANCD2/FANCI complex is tightly interconnected (27), we would expect mUb-FANCI to be similarly affected by UL138 but lack the reagents to measure it directly.

**Figure 4.**
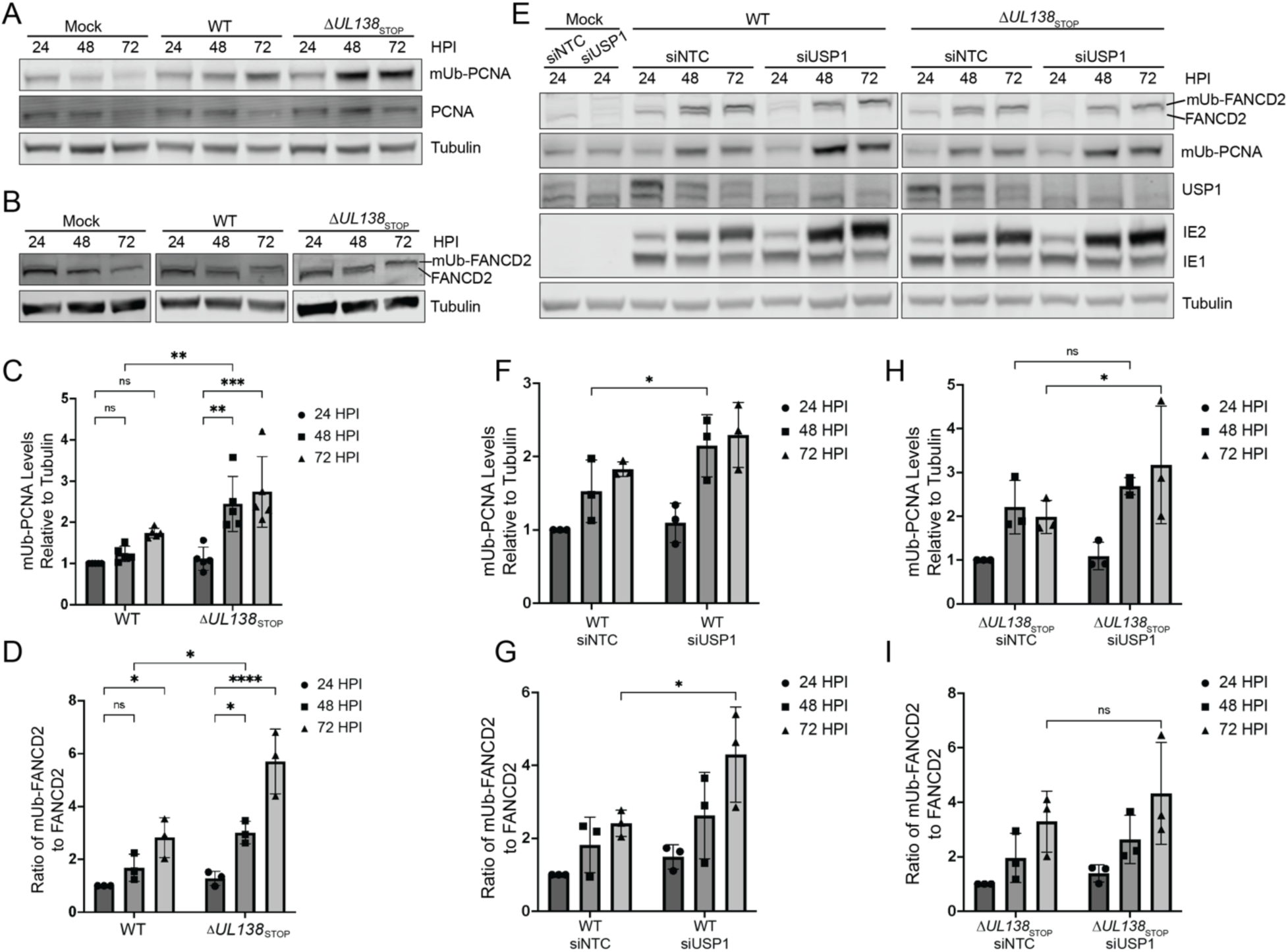
UL138 and USP1 drive deubiquitination of PCNA and FANCD2 in infection. (A-D) Fibroblasts were mock-infected or infected (MOI = 1) with TB40/E-WT or -Δ*UL138*_STOP_ virus over a 72-hour time course. Immunoblotting was performed on whole cell lysates collected at the indicated time points. Tubulin serves as a loading control. (A) PCNA and mUb-PCNA were detected using monoclonal antibodies specific to each and secondary antibodies conjugated to DyLight™ 680 (mouse) or 800 (rabbit). (B) FANCD2 and mUb-FANCD2 were detected using a polyclonal antibody to FANCD2 and a secondary antibody conjugated to DyLight™ 800. (C) Quantification of mUb-PCNA relative to tubulin at each time point and normalized to WT at 24 hpi. (D) Quantification of the ratio of mUb-FANCD2 to FANCD2 normalized to WT at 24 hpi. (E) Fibroblasts were reverse transfected with a siRNA pool targeting USP1 or a NTC. At two days post transfection, cells were mock-infected or infected with WT or Δ*UL138*_STOP_ virus, and whole cell lysates were collected over a 72-hour time course. IE1 and IE2 are HCMV immediate early proteins used as controls for infection. (F-G) Quantification of mUb-PCNA relative to tubulin (F) and the ratio of mUb-FANCD2 to FANCD2 (G) in WT infection normalized to the siNTC at 24 hpi. (H-I) Quantification of mUb-PCNA relative to tubulin (H) and the ratio of mUb-FANCD2 to FANCD2 (I) in Δ*UL138*_STOP_ infection relative to the siNTC at 24 hpi. For all statistical analyses, significance was determined by two-way ANOVA with Tukey’s multiple comparisons test. Asterisks (*P<0.05, **P<0.01, ***P<0.001, ****P<0.0001) represent statistically significant differences determined in a minimum of three independent experiments.

To confirm that deubiquitination of PCNA and FANCD2 was mediated through USP1 in virus infection, we depleted USP1 using siRNA knockdown. Compared to the NTC, USP1 depletion in WT infection led to a modest, but significant, increase in mUb-PCNA and mUb-FANCD2 at 48 hpi and 72 hpi, respectively, resembling the phenotype observed in Δ*UL138*_STOP_ infection (Fig. 4E). Therefore, USP1 also mediates deubiquitination of these proteins in WT CMV infection. USP1 depletion in Δ*UL138*_STOP_ infection (Fig. 4E) increased mUb-PCNA (Fig. 4H), but did not have a statistically significant effect on mUb-FANCD2 (Fig. 4I) compared to the NTC control. Taken together, these data support the hypothesis that UL138 drives USP1 complex activity to deubiquitinate mUb-PCNA and mUb-FANCD2 in CMV infection.

### USP1-regulated DNA repair pathways restrict CMV replication

We next sought to determine the overall significance of USP1-regulated DNA repair pathways during CMV replication. Previously we found that depletion of PCNA resulted in increased vDNA synthesis and virus replication, suggesting PCNA alone restricts CMV TB40/E replication (19). We also showed that this restriction was dependent on the mUb of PCNA. To assess how USP1 DNA repair substrates influence CMV replication, we depleted PCNA, FANCD2, and FANCI [si(PFF)] in combination and with additional knockdown of USP1 [si(U-PFF)] and then measured viral genome copy number and virus yields. siRNA knockdown resulted in ≥50% depletion of USP1, PCNA, and FANCD2 (Fig. 5A). While we could not verify knockdown of FANCI protein, depletion of FANCD2 is presumed to impact function of the FANCD2/FANCI complex (27). Compared to NTC, depletion of PCNA/FANCD2/FANCI or PCNA/FANCD2/FANCI combined with USP1 increased viral genome synthesis in WT and Δ*UL138*_STOP_ infections (Fig. 5B). Consistent with this, we also observed that depletion of these factors increased virus yield in WT infection (Fig. 5C). Similar to findings with USP1 knockdown alone, Δ*UL138*_STOP_ yields also trended up with si(PFF) or si(U-PFF), but this increase did not reach statistical significance. Therefore, USP1-PCNA/FANCD2/FANCI DNA repair system restricts CMV vDNA synthesis and virus replication in a manner modulated by UL138.

**Figure 5.**
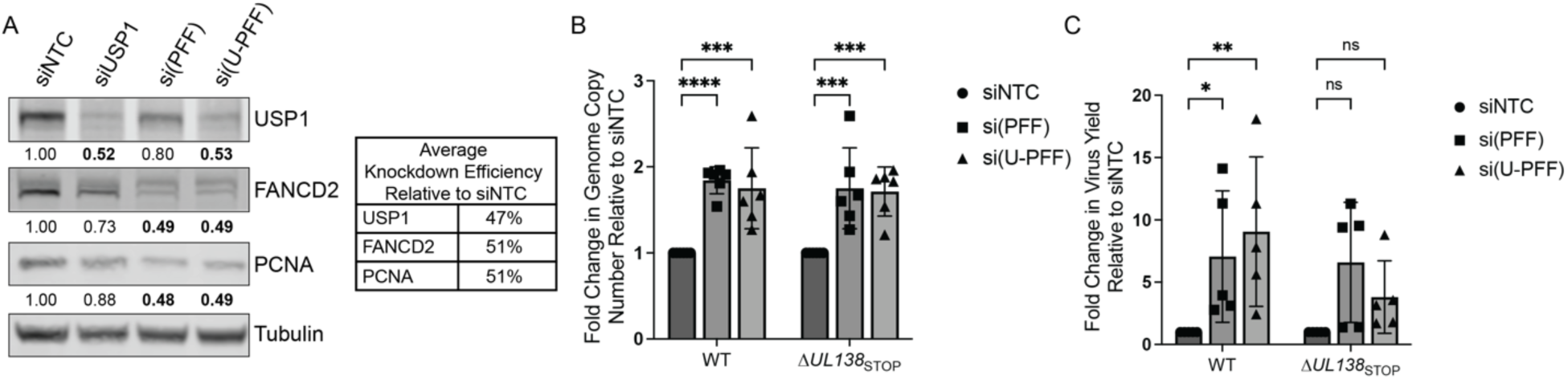
USP1-regulated DNA repair pathways restrict HCMV replication. Fibroblasts were reverse transfected with a siRNA pool targeting USP1, PCNA, FANCD2, FANCI, or a nontargeting control (NTC). For pooled knockdowns, si(PFF) represents combined knockdown of PCNA, FANCD2, and FANCI, and si(U-PFF) includes USP1 in addition to PCNA/FANCD2/FANCI. (A) At two days post transfection, whole cell lysates were collected for immunoblotting. Indicated proteins were detected using primary antibodies and secondary antibodies conjugated to DyLight™ 680 (mouse) or 800 (rabbit). Band density was quantified relative to tubulin and normalized to siNTC to determine average knockdown efficiency. (B-C) At two days post transfection, cells were infected (MOI = 1) with TB40/E-WT or -Δ*UL138*_STOP_ and collected at 96 hpi. (B) Viral genome copy number was determined by qPCR using a TB40/E BAC standard curve and primer set designed for the region of the viral genome encoding the β2.7 transcript. The graph represents the fold change in viral genome copy number relative to siNTC. (C) Virus yields were measured by TCID_50_ and normalized relative to siNTC. Statistical significance was determined by two-way ANOVA with Dunnett’s multiple comparisons test. Asterisks (*P <0.05, **P <0.01, ***P <0.001, ****P <0.0001) represent statistically significant differences determined for four independent experiments.

### UL138 and USP1-PCNA/FANCD2/FANCI contribute to CMV genome integrity

This work demonstrates a role for UL138 in regulating host DNA synthesis and repair pathways through its interaction with USP1 with consequences for vDNA synthesis and replication. We previously demonstrated that host TLS polymerases were recruited to viral RCs and functioned to maintain CMV genome integrity (13). As USP1 regulates the recruitment of TLS polymerases through mUb-PCNA(22,23), we hypothesized a role for UL138-USP1 and the modulation of PCNA/FANCD2/FANCI ubiquitination in regulating viral genome integrity. To explore this, we investigated how depletion of these host factors impacts stability of the viral genome. USP1, PCNA/FANCD2/FANCI, or all four factors together were depleted in growth-arrested cells and cells were infected with WT or Δ*UL138*_STOP_ virus. At 96 hpi, we extracted DNA from infected cells for Illumina sequencing. All sequence reads mapping to the human genome were removed. The remaining reads were aligned to the TB40/E reference genome. Structural variants present in the TB40/E-WT virus inoculum were subtracted from all reads to exclude pre-existing rearrangements present in the WT virus stock. This approach was taken (rather than subtracting WT and Δ*UL138*_STOP_ condition from its respective inoculum) to gain insight into differences in rearrangements arising purely from the lack of UL138, in addition to the absence of USP1 or USP1-PCNA/FANCD2/FANCI. We quantified structural variants indicated by novel DNA junctions created by duplications, inversions, and deletions that arose during virus replication under each infection condition (13). We achieved a minimum sequencing depth of 100x per sample and focused our analysis on novel DNA junctions that occurred with a minimum frequency of 2.5% across five biological replicates. Circular renditions of the genome are shown (Fig. 6 A-B) with the frequency of novel junctions represented by arcs spanning the junctions.

**Figure 6.**
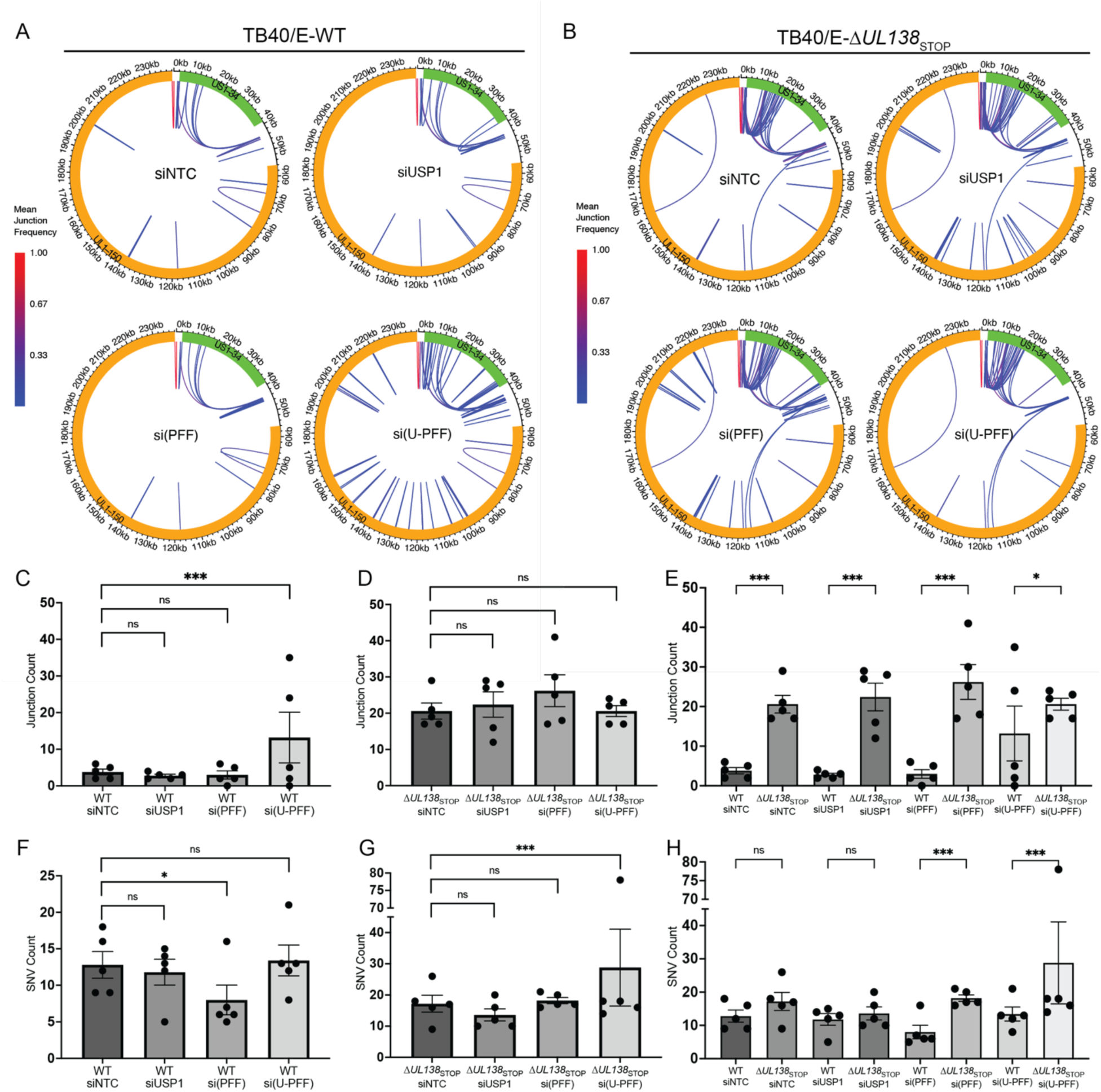
UL138 and USP1-PCNA/FANCD2/FANCI promote CMV genome integrity through. Fibroblasts were reverse transfected with a siRNA pool targeting USP1, PCNA/FANCD2/ FANCI (si-(PFF)), a combination of all 4 (si(U-PFF)) or a nontargeting control (NTC). At two days post transfection, cells were infected (MOI = 1) with TB40/E-WT or - Δ*UL138*_STOP_ and total DNA was isolated at 96 hpi for sequencing. Sequences from each knockdown condition, as well as from the virus stocks used for infection, were aligned to the TB40/E-GFP reference genome. Structural variants present in the TB40/E-WT virus inoculum were subtracted from all reads to exclude pre-existing rearrangements present in the WT virus stock. (A-B) Mean novel junction (inversions, deletions, duplications) frequency within each condition for (A) WT and (B) Δ*UL138*_STOP_ infections are shown. HCMV genomic coordinates are plotted along the circular axis and the UL (orange) and US (green) regions of the genome are marked. The arcs connect novel junction points detected at the average frequency for the given condition indicated by the color scale. (C-E) Quantification of the number of novel junctions detected per sample (n = 5) for each condition: (C) WT, (D) Δ*UL138*_STOP_ infection, and (E) pairwise quantifications of both infections. (F-H) Quantification of the number of single nucleotide variations (SNVs: point mutations, insertions, deletions) detected per sample (n = 5) for each condition: (F) WT, (G) Δ*UL138*_STOP_ infection, and (H) pairwise quantifications of both infections. Statistical significance was determined by pairwise two-sided exact Poisson tests and adjusted using Bonferroni correction (*P <0.05, **P <0.01, ***P <0.001).

Within WT-infected cells, we found that knockdown of USP1 or PCNA/FANCD2/FANCI alone was not sufficient to result in significant changes in genomic rearrangements compared to the siNTC condition (Fig. 6A, quantified in 6C). However, knockdown of USP1 in combination with PCNA/FANCD2/FANCI, si(U-PFF), increased genomic rearrangements relative to siNTC. While this was driven by two replicates, our observations align with the stochastic nature of DNA rearrangement events, which may be influenced by the architectural features of the viral genome or rate of replication (51). These results indicate a role for USP1-regulated DNA repair pathways in maintaining viral genome integrity. Relative to the Δ*UL138*_STOP_-NTC condition, depletion of USP1, PCNA/FANCD2/FANCI, or the combination did not trigger additional increase in novel junctions relative to the junctions created by the loss of *UL138* alone (Fig. 6B, quantified in 6D). This result suggests a role for UL138 in directing the action of USP1-PCNA/FANCD2/FANCI to ensure viral genome integrity.

Notably, loss of UL138 alone increased structural variants across the viral genome compared to WT infection regardless of knockdown condition (Fig. 6E). This includes novel junctions that span large regions in the unique long (UL) region that have not been observed in TLS (13), PCNA(19), or USP1-PCNA/FANCD2/FANCI knockdowns, suggesting additional roles for UL138 in maintaining viral genome integrity during productive infection independent of USP1-PCNA/FANCD2/FANCI. Further, Δ*UL138*_STOP_-associated junctions differ in distribution across the genome relative to those arising due to depletion of USP1-PCNA/FANCD2/FANCI in WT infection (Fig. 6A, si(U-PFF)), suggesting qualitative differences in the junctions arising in each condition. The number of sequencing reads was equalized across all treatments by subsampling (13), and therefore increased junction counts in the mutant virus cannot be explained by a simple increase in number of genomes per cell. These results highlight UL138 as a key player in protecting CMV genome integrity.

We previously found that PCNA and TLS polymerases ρι, κ, and 1 contribute to CMV genome diversity where depletion of these host factors decreased SNVs (point mutations, insertions, or deletions) in the viral genome (13,19). Consistent with this result, depletion of PCNA/FANCD2/FANCI in WT infection resulted in decreased SNVs on vDNA (Fig. 6F). This effect was lost with the additional knockdown of USP1. This result is suggestive of the role of monoubiquitination in regulating TLS-generated SNVs. We would anticipate residual PCNA/FANCD2/FANCI remaining due to incomplete knockdown to be more heavily monoubiquitinated in the absence of USP1, possibly heightening TLS recruitment, relative to where USP1 is present (15). By contrast, in Δ*UL138*_STOP_ infection SNV counts increased in the USP1-PCNA/FANCD2/FANCI knockdown relative to any other knockdown in Δ*UL138*_STOP_ infection (Fig. 6G and H), although this was driven by a single replicate. These results suggest a minimal impact of UL138 on pathways driving SNVs.

Taken together, these results demonstrate an important role for USP1-PCNA/FAND2/FANCI and UL138 in maintaining viral genome integrity. The finding that some structural variants (e.g., those spanning large regions in the UL region) are unique to the loss of UL138 and not observed in USP1-PCNA/FANCD2/FANCI knockdowns suggests that UL138 has roles in regulating distinct DDR pathways, some of which are independent of the roles of USP1-PCNA/FANCD2/FANCI. These results further suggest a more substantial role for the UL138-USP1-PCNA/FANCD2/FANCI interaction in regulating DNA pathways to prevent large structural rearrangements rather than SNVs.

### UL138 directs USP1-dependent host HDR pathways

To gain more insight into the role of UL138 and USP1-PCNA/FANCD2/FANCI in controlling DDR pathways and maintaining CMV genome integrity, we analyzed the distribution and type (e.g., inversions, deletions, or duplications) of structural variants in each of our infection and knockdown conditions (Fig. 7). Δ*UL138*_STOP_ infection increased structural variants across the CMV genome (0-236 kb) relative to WT infection, as indicated by the increase in novel junctions (Fig. 6E), these occurred across both the unique short (US, 0-56 kb) region and the unique long (UL, 56-236 kb) regions (Fig. 7A). When we normalize junction counts relative to the size of the UL and US genomic regions, the susceptibility and prominence of structural variants in the US region is more apparent (Fig. 7B). Inversions were the predominant structural variant in all WT infections (Fig. 7C). While inversions also predominated in Δ*UL138*_STOP_ infections (Fig. 7C), the loss of UL138 increased deletions relative to WT infection (Fig 7C). However, Δ*UL138*_STOP_ infection exhibited an increase in all types of junctions relative to WT infection (Fig. 7D). Novel junctions in both the WT and Δ*UL138*_STOP_ infection predominantly accumulated in the 0-56 kb region (Fig. 7E), although the loss of UL138 also resulted in accumulation of inversions in the 56-236 kb region.

**Figure 7.**
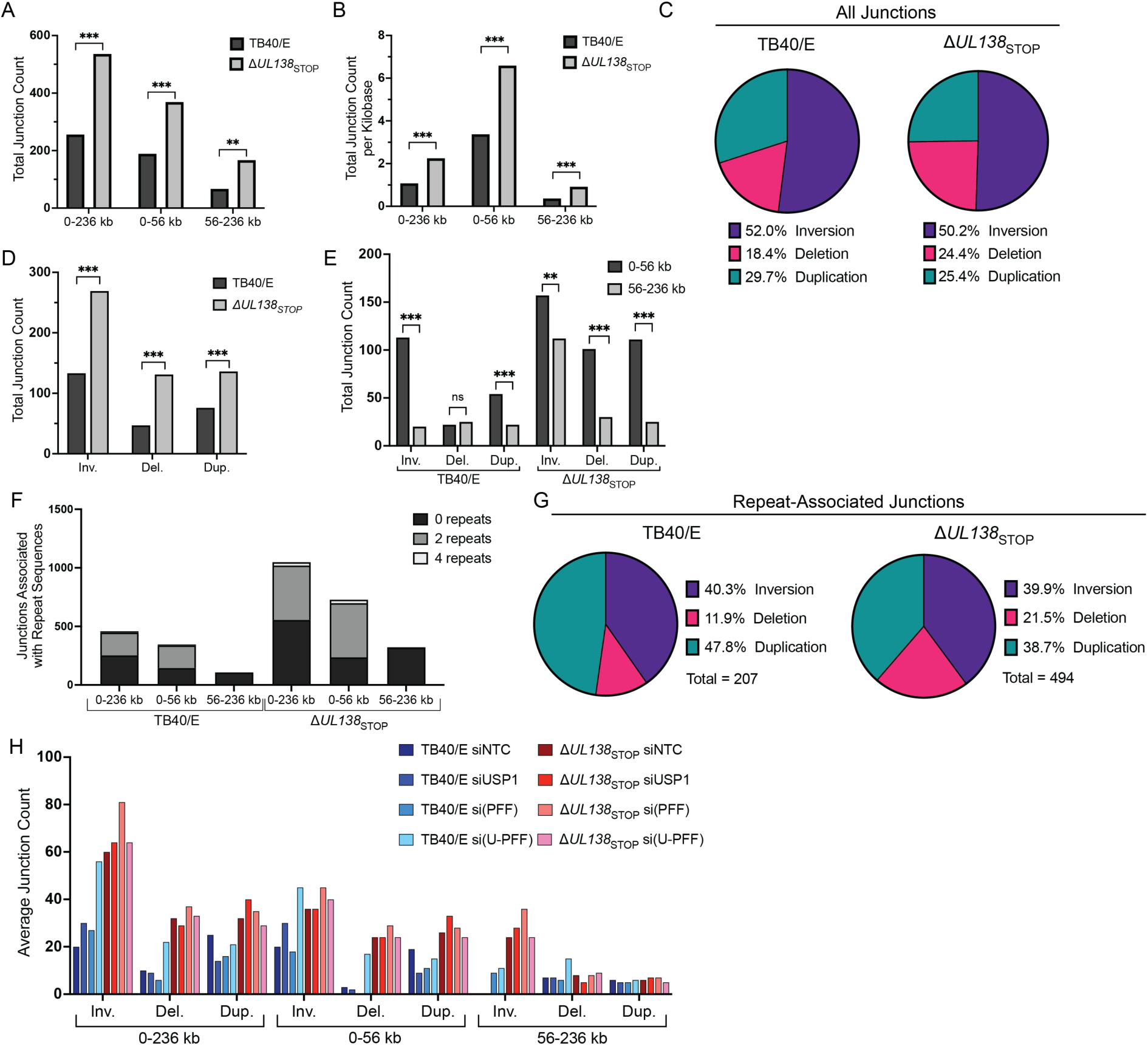
UL138 directs USP1-dependent and -independent host DDR pathways. (A) Total number of novel junctions in TB40/E-WT and -Δ*UL138*_STOP_ infections separated by region of genome in which they occur: 0-236 kb (whole genome), 0-56 kb (US region), 56-236 kb (UL region). Statistical analysis: exact Poisson test. (B) Total number of novel junctions as a proportion of the number of kilobases in each genomic region. Statistical analysis: Wilcoxon rank sum test. (C) Type of structural variant (inversion, deletion, duplication) as a percentage of all novel junctions in both infections. (D) Total number of novel junctions across the whole genome, separated by junction type, for WT and Δ*UL138*_STOP_ infections. Statistical analysis: exact Poisson test. (E) Total number of novel junctions in each genomic region, separated by junction type for both infections. Statistical analysis: exact Poisson test. (F) Quantification of novel DNA junctions associated with repeat sequences in the CMV genome for WT and Δ*UL138*_STOP_ infections. Repeats are defined as 20-nucleotide sequences on either side of the novel junction found in the reference genome with 100% identity. “0 repeats” represents the 20-nucleotide sequences flanking the junction were unique. (G) Type of junction as a percentage of all repeat-associated junctions occurring in WT and Δ*UL138*_STOP_ infections. (H) Average number of novel junctions in each genomic region separated by junction type for all knockdown conditions within WT and Δ*UL138*_STOP_ infections. For all statistical analyses, ***P <0.001, **P <0.01, *P <0.05.

We previously observed that structural variants arising from depletion of TLS polymerases were associated with repeat sequences in the CMV genome (13). Therefore, we examined junctions associated with 20-nucleotide repeating sequences across the viral genome. While loss of UL138 increased structural variants overall, it also resulted in increased structural variants associated with sequences that are repeated 2 or more times, particularly in the 0-56 kb of the genome (Fig. 7F). Notably, junctions arising in the 56-236 kb region were not associated with repeats in either WT or Δ*UL138*_STOP_ infection. Further, more repeat-associated deletions increased in Δ*UL138*_STOP_ infection compared to WT (Fig. 7G). Similar to the knockdown of TLS polymerases (13), most junctions were associated with GC-rich regions of the entire genome across all infections (Fig. S2), and this association was not demonstratively altered between WT and Δ*UL138*_STOP_ infections.

To better understand the specific contributions of USP1-PCNA/FANCD2/FANCI and its modulation by UL138 in protecting against specific types of structural variants in the US or UL regions of the viral genome, we further parsed out the types of structural variants formed by each knockdown/infection condition (Fig. 7H). The combined depletion of USP1-PCNA/FANCD2/FANCI in WT infection triggered a robust increase in inversions and deletions across the entire CMV genome (Fig. 7H, lightest blue bars) relative to the siNTC. While inversions and deletions were increased in Δ*UL138*_STOP_ infections relative to the WT infection, there was no additional impact of knockdowns in Δ*UL138*_STOP_ infections (Fig. 7H, red bars).

This suggests that UL138 protects against inversions and deletions in a manner dependent on USP1-PCNA/FANCD2/FANCI. However, we note qualitative differences in the location of these rearrangements as they were primarily constrained to the US region (0-56 kb), which is associated with repeat sequences (Fig. 7F and H). In the UL region (56-236 kb), inversions were associated with loss of UL138 and unaffected by the loss of USP1-PCNA/FANCD2/FANCI in either WT or Δ*UL138*_STOP_ infection and were not associated with repeat sequences (Fig. 7F and H). As inversions and deletions associated with repeat sequences are a hallmark of HDR, these data are consistent with the hypothesis that UL138 and USP1-PCNA/FANCD2/FANCI protects genome integrity through HDR pathways. These results also suggest that UL138 regulates distinct (non-USP1-PCNA/FANCD2/FANCI-driven and non-repeat driven) DDR pathways that remain to be defined.

As the loss of UL138 alone results in structural variants, we suspected that structural variants in Δ*UL138*_STOP_ infections might be present in the stocks used for infection. Therefore, we compared the sequences from the viral stocks to the TB40/E reference genome to determine the extent of variations accumulating during Δ*UL138*_STOP_ stock preparation. The Δ*UL138*_STOP_ stock virus contained more structural variants relative to the reference genome than the WT virus (Fig. 8A). The most striking change is the increase of deletions in the US region (also apparent in the data set from infection, Fig. 7C). These data suggest an inherent instability due to the loss of UL138. It is important to note that we derive virus stocks from only one round of replication after reconstitution of virus from the bacterial artificial chromosome (BAC) and we never serially passage virus in the preparation of stocks. To ensure that the deletions that accumulated during Δ*UL138*_STOP_ replication were not the result of an aberrant BAC clone from which virus stocks are derived, we sequenced the BAC DNA used for the initial propagation of TB40/E-WT and -Δ*UL138*_STOP_ virus stocks. We observed no major changes in the BAC sequences compared to the TB40/E reference (Fig. 8B-C), validating that the instability of the viral genome in the absence of UL138 is not an artifact derived from BAC recombineering.

**Figure 8.**
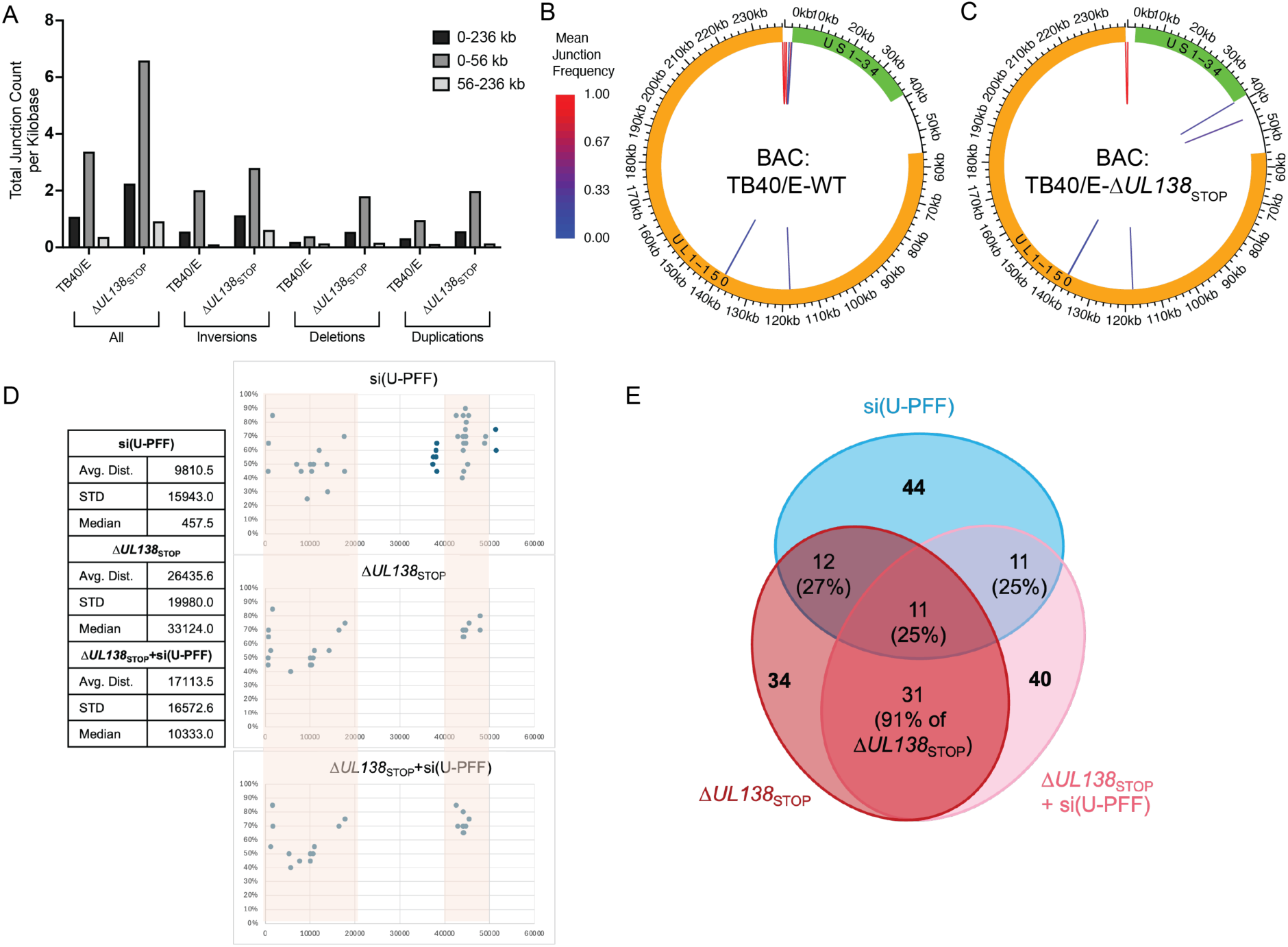
Analysis of novel junction origins. (A) Total number of junctions per kilobase of genomic region for all junctions and distinct junction types in TB40-E-WT and -Δ*UL138*_STOP_ virus stocks used for infection. All junctions are shown on the left and then separated into junction type. (B-C) Bacterial artificial chromosomes containing the TB40/E-WT or -Δ*UL138*_STOP_ genome were purified and sequenced. Sequences were subsequently aligned to the TB40/E reference genome. Mean novel junction (inversions, deletions, duplications) frequency within each condition for (B) WT and (C) Δ*UL138*_STOP_ alignments are shown. HCMV genomic coordinates are plotted along the circular axis and the UL (orange) and US (green) regions of the genome are marked. The arcs connect novel junction points detected at the average frequency for the given condition. (D) The average distance, standard deviation, and median distance between inversions in the US region was calculated for the conditions shown (see table). Junctions are plotted across the US regions with the region containing overlapping junctions shaded. (E) Overlap of inversions arising in the three conditions in panel D are shown. Total number of junctions are shown in bold font and the proportion overlapping indicated.

Finally, we were intrigued by finding that depletion of USP1-PCNA/FANCD2/FANCI did not increase junctions beyond the disruption of *UL138* in infection (Fig. 7H). This is consistent with UL138 and USP1-PCNA/FANCD2/FANCI working in the same pathway, as we would hypothesize from their interaction. To explore this further, we calculated the average distance, standard deviation and median distance between inversions in the US (0-56kb) region as the most prominent novel junctions. While depletion of USP1-PCNA/FANCD2/FANCI produced some unique rearrangements, most were clustered in the first 20-kbp or within 40-50-kbp of the US region (Fig. 8D). Strikingly, the junctions arising due to the disruption of *UL138* (Δ*UL138*_STOP_) or combined loss of *UL138* and USP1-PCNA/FANCD2/FANCI were highly concordant with 91% overlap (Fig. 8D-E). Only 25% of the novel junctions arising from depletion of USP1-PCNA/FANCD2/FANCI overlap with Δ*UL138*_STOP_ infections, suggesting an important role for UL138 in directing these pathways. An alternative explanation is the high overlap in junctions is due to a founder effect from junctions already present in the virus inoculum. However, when comparing junctions present in the Δ*UL138*_STOP_ virus stock to junctions arising through replication in each of the Δ*UL138*_STOP_ infections, we find that 47% (16/34) or 45% (18/40) of junctions present in the inoculum were in the Δ*UL138*_STOP_ siNTC or Δ*UL138*_STOP_ si(U-PFF) infections. Therefore, founder effects cannot account for the majority of the identical junctions seen in Δ*UL138*_STOP_ siNTC and Δ*UL138*_STOP_ si(U-PFF) infections. These results further support our finding that the interaction between UL138 and USP1 functions to direct PCNA/FANCD2/FANCI mediated repair pathways in HCMV infection.

## DISCUSSION

DNA replication coordinates a myriad of proteins to synthesize DNA while simultaneously sensing and responding to damage. For example, resolution of an interstrand crosslink may require interplay of multiple pathways—FA, TLS, HDR and nucleotide excision repair (52)—all in order to maintain integrity of the genome. Viral infection introduces foreign genetic material and proteins into the host cell, presenting a potential challenge to host genome integrity. Many viruses, including CMV, induce a DNA damage response upon infection, which could simply represent a host defense against virus replication (11). However, viruses often exploit cellular responses, utilizing host proteins for vDNA synthesis and, in the case of viruses that coexist with their host, also use host factors to protect viral genome maintenance (2–4,53,54). These pathways may also drive viral diversity. The full extent to which CMV hijacks the host DDR is not yet defined and these likely impact viral DNA synthesis, as well as genome maintenance in latency. This work defines the role of a viral protein, UL138, and a host pathway it targets in regulating interconnected host DNA repair pathways, which contribute to virus replication and genome integrity. Furthermore, because the viral genome does not integrate into the host chromosomes this work introduces CMV as a powerful exogenous model-chromosome for understanding complex interplay between host pathways, as the virus infects and replicates its genome in growth-arrested primary cells that tolerate the knockdown of essential host factors.

USP1 constrains CMV genome synthesis and overall replication during productive infection of fibroblasts (Fig. 1B), paralleling the observation that chemical inhibition of USP1 stimulates virus replication in hematopoietic progenitor cells where viral latency is established (28). The UAF1-USP1 complex has also been reported to facilitate genome replication of human papillomavirus (HPV) through interaction with the viral-encoded E1 helicase (55). In addition to DNA replication and repair, UAF1-USP1 has been reported to regulate antiviral responses (56,57), and we have shown that UL138-USP1 upregulates STAT1 innate immune signaling (28,56,57). Moreover, it was recently shown that herpes simplex virus (HSV-1) induces the TLS pathway early during infection but the viral-encoded deubiquitinating enzyme, UL36USP, counters this to evade innate immune activation through cGAS (58). The UL138-USP1 interaction suggests a means by which CMV can coordinate the regulation of DDR and innate signaling pathways, and future work will be aimed at differentiating the roles between UAF1-USP complexes in CMV infection, as well as understanding the relationship between innate immune sensing and DDR pathways.

Notably, UAF1 is required for efficient CMV replication (Fig. 1). This finding is consistent with previous studies using the Han strain of CMV (38) and suggests that other UAF1-USP complexes play a significant role in virus infection. Both USP12 and USP46 regulate signaling in the AKT pathway (59,60), which is also important to CMV replication and latency (61–63). AKT signaling is downregulated during productive infection of fibroblasts, and inhibition of AKT or its downstream pathways is a potent stimulator of CMV reactivation in hematopoietic progenitor cells (61,62). Beyond AKT, USP12 also stimulates STAT1 signaling (64) and USP46 regulates the cell cycle through stabilizing the CRL4^Cdt2^ E3 ubiquitin ligase complex, which promotes proliferation of HPV-transformed cancers (65). Interestingly, the USP46-CRL4^Cdt2^ interaction also has implications for genome integrity as it regulates DNA repair proteins, including the TLS polρι, dependent on its interaction with PCNA (66). Therefore, depletion of UAF1 has pleiotropic effects at the innate-DDR signaling nexus that remain to be understood.

The re-localization of USP1, PCNA, and FANCD2 to viral RCs suggests that these host proteins function in viral DNA synthesis. We found that UL138 regulates USP1-mediated deubiquitination of PCNA and FANCD2, a modification that impacts the function of these proteins in DNA repair (20,27). Beyond PCNA and FANCD2, USP1 has additional nuclear substrates, including RAD51 associated protein 1 (RAD51AP1) (67) and inhibitors of DNA binding (ID) proteins (68). USP1 promotes DNA repair by HDR through interaction with RAD51AP1 (67). USP1 deubiquitination and stabilization of ID proteins prevents osteosarcoma cell differentiation (68). These will be important targets to investigate going forward in understanding the role of UL138 in modulating DDR pathways in the context of CMV infection. These interactions may contribute to oncomodulatory features of CMV.

The FA repair pathway has also been reported to be activated upon infection of other viruses. The E6 and E7 oncoproteins of high-risk HPV increase mUb-FANCD2 while E6 disrupts its deubiquitination by USP1 (66). FA repair is also activated in simian virus 40 (SV40) infection and required for efficient virus replication (69), but specific contributions to infection remain unknown. Adenovirus (Ad5) replication induces the FA pathway, and FANCD2 is required for optimal Ad5 replication as a contributor to replication-dependent recombination (70). Similar to findings presented here, HSV-1 productive infection is associated with increased mUb-FANCD2 which is also re-localized to RCs (71). Karttunen and colleagues report that FANCD2 is required for HSV-1 infection and that activation of the FA pathway suppresses non-homologous end joining (NHEJ), an important step in the HSV-1 life cycle (71). CMV infection induces re-localization of various NHEJ proteins and alters total protein levels (12), but their contribution to infection is uncharacterized. While further work is required to define specific roles of FANCD2 in CMV infection, our data suggests the FA pathway is important for repair of the viral genome during replication.

UL138 is a protein that primarily localizes to Golgi compartments (45) and has not been reported to have nuclear functions during productive infection. It is possible that UL138 modulates the activity of the UAF1-USP1 complex in the cytoplasm prior to nuclear trafficking. In support of this, we observed that USP1 co-localized with UL138 in the Golgi apparatus during infection (Fig. S3A). However, UL138 had no effect on total levels of USP1 (Fig. S3B-C) nor its localization within RCs (Fig. S1) despite significant differences observed with ubiquitination of USP1 substrates and viral genomic rearrangements. Future work will be aimed at defining a specific mechanism for these effects.

We previously observed that PCNA restricted CMV replication in a manner dependent on modification at lysine 164, the site of mUb and SUMOylation (19). Our findings here support those observations whereby depletion of PCNA in conjunction with FANCD2/FANCI relieved a restriction to vDNA synthesis (Fig. 5). Given that PCNA alone, in contrast to TLS polymerases, was not required for viral genome integrity (19), it is possible that the restriction of vDNA synthesis results from an alternative (non-canonical bypass/TLS) form of repair. While non-canonical roles of TLS polymerases are poorly understood, CMV offers a powerful model going forward to investigate these functions. Further, depletion of USP1-PCNA/FANCD2/FANCI was sufficient to compromise viral genome integrity in WT infection (Fig. 6). The increase in structural rearrangements across the genome, particularly those associated with repeat sequences, with depletion of USP1-PCNA/FANCD2/FANCI pathway is a signature of dysregulated HDR pathways.

There was a striking mutational signature associated with the loss of UL138 in infection. Loss of UL138 increased CMV genomic instability, as evidenced by an overall increase in structural rearrangements. The increased rearrangements present in *UL138*-mutant virus infection could result from the absence of a pathway UL138 recruits or the intervention of an alternative repair pathway recruited in the absence of UL138. Further, the data suggests UL138 regulates multiple pathways in each region of the genome. UL138 and USP1-PCNA/FANCD2/FANCI were important for protecting viral genome integrity in the US (0-56 kb). The rearrangements derived from the loss of UL138 and loss of UL138 combined with the depletion of USP1-PCNA/FANCD2/FANCI were largely the same (91% overlap) suggesting one pathway modulated by UL138 with USP1-PCNA/FANCD2/FANCI. By contrast UL138 was important for integrity of the UL (56-236 kb) region independently of USP1-PCNA/FANCD2/FANCI. The novel junctions arising in the US region of the CMV genome has an increased association with repeat sequences, suggesting a role for UL138-USP1-PCNA/FANCD2/FANCI in modulating an HDR pathway. By contrast, the rearrangements in the UL region arising in *UL138*-mutant infection occurred independently of repeat sequences and USP1-PCNA/FANCD2/FANCI. This suggests that fundamentally different host pathways act on the US and UL regions of the CMV genome. While the deletions and inversions associated with repeat sequences are indicative of HDR, non-repeat associated inversions arise through repair mechanisms including, NHEJ, break induced replication or template switching (51). While, these observations implicate a role for UL138 in regulating double-strand break repair pathways (72), perhaps by modulating activity of factors that determine which repair pathways prevail at any given DNA break, further work is required to define specific repair mechanisms by which these viral and host proteins protect against structural rearrangements to ensure viral genome integrity.

We have demonstrated a role for USP1 and its substrates in maintaining integrity of the CMV genome during replication, revealing novel insights into a long-standing question as to how this axis of DDR induced by DNA virus infection contributes to viral genome synthesis. We also identify the first CMV protein that modulates this pathway, as well as an additional pathway that remains to be defined, and actively contributes to the integrity of viral genomes. While UL138 is required to maintain genome integrity in both the US and UL regions, this occurs through distinct pathways, with USP1-PCNA/FANCD2/FANCI acting exclusively on repeat associated sequences in the US region. Moving forward, it will be important to understand mechanisms by which UL138 and USP1 direct HDR and other repair pathways. The change in SNVs depending on the presence of PCNA/FANCD2/FANCI and UL138 indicates that lesion bypass occurs on the HCMV genome (Fig. 6F-H) (13) and suggests a role for TLS polymerases. However, our results indicate additional roles for UL138 and USP1-PCNA/FANCD2/FANCI in regulating HDR or break-induced repair pathways. It will be of interest to determine the role for TLS polymerases in these pathways. HCMV, as a complex model DNA genome, serves as a powerful extrachromosomal tool to investigate DDR pathways in primary human cells. Further, this work has important implications for maintenance of the viral genome for latency, which is the focus of ongoing work.

## DATA AVAILABILITY

Alignments used for Figure 6 are available at the University of Arizona Research Data Repository (doi: 10.25422/azu.data.28383527). Raw sequence reads have been deposited in the Sequence Read Archive, https://www.ncbi.nlm.nih.gov/sra (BioProject ID: PRJNA1226608).

## ACKNOWLEDGMENTS

We are grateful to Dr. James Alwine (University of Pennsylvania), and Dr. Lynn Enquist (Princeton University) for critical discussion. We acknowledge Dr. Ross Buchan (University of Arizona) for assistance with deconvolution microscopy.

## AUTHOR CONTRIBUTIONS

Pierce Longmire: Conceptualization, Data curation, Formal analysis, Investigation, Methodology, Resources, Visualization, Writing – original draft, Writing – review and editing. Sebastian Zeltzer: Conceptualization, Data curation, Investigation, Methodology, Supervision, Writing – review and editing. Kristen Zarrella: Conceptualization, Data curation, Investigation, Methodology, Writing – review and editing. Olivia Daigle: Formal analysis, Investigation, Methodology, Visualization, Writing – review and editing. Marek Svoboda: Data curation, Formal analysis, Methodology, Writing – review and editing. Justin M. Reitsma: Methodology, Writing – review & editing. Scott S. Terhune: Data curation, Writing – review and editing. Carly Bobak: Conceptualization, Data curation, Formal analysis, Investigation, Supervision, Writing – review and editing. Giovanni Bosco: Conceptualization, Data curation, Formal analysis, Investigation, Funding acquisition, Project administration, Supervision, Writing – review and editing. Felicia Goodrum: Conceptualization, Formal analysis, Funding acquisition, Project administration, Supervision, Writing – original draft, Writing – review and editing.

## REFERENCES

1. Weitzman, M.D., Lilley, C.E. and Chaurushiya, M.S. (2010) Genomes in Conflict: Maintaining genome integrity during virus infection. Annual review of microbiology, 64, 61–81.

2. Weller, S.K. and Sawitzke, J.A. (2014) Recombination promoted by DNA viruses: phage λ to herpes simplex virus. Annual review of microbiology, 68, 237–258.

3. Luftig, M.A. (2014) Viruses and the DNA Damage Response: Activation and Antagonism. Annual review of virology, 1, 605–625.

4. McFadden, K., Luftig, M.A., Cullen, B.R. and Cullen, B.R. (2013) Interplay Between DNA Tumor Viruses and the Host DNA Damage Response. CURR TOP MICROBIOL, 371, 229–257.

5. Li, S., Liu, B., Tan, M., Juillard, F., Szymula, A., Alvarez, A.L., Van Sciver, N., George, A., Ramachandran, A., Raina, K., et al. (2024) Kaposi’s sarcoma herpesvirus exploits the DNA damage response to circularize its genome. Nucleic acids research, 52, 1814–1829.

6. Dolan, A. (2004) Genetic content of wild-type human cytomegalovirus. J Gen Virol, 85, 1301–1312.

7. Murphy, E., Yu, D., Grimwood, J., Schmutz, J., Dickson, M., Jarvis, M.A., Hahn, G., Nelson, J.A., Myers, R.M. and Shenk, T.E. (2003) Coding Potential of Laboratory and Clinical Strains of Human Cytomegalovirus. Proceedings of the National Academy of Sciences - PNAS, 100, 14976–14981.

8. Murphy, E., Rigoutsos, I., Shibuya, T. and Shenk, T.E. (2003) Reevaluation of Human Cytomegalovirus Coding Potential. Proceedings of the National Academy of Sciences - PNAS, 100, 13585–13590.

9. Ravichandran, S., Kim, Y.-E., Bansal, V., Ghosh, A., Hur, J., Subramani, V.K., Pradhan, S., Lee, M.K., Kim, K.K. and Ahn, J.-H. (2018) Genome-wide analysis of regulatory G-quadruplexes affecting gene expression in human cytomegalovirus. PLoS pathogens, 14, e1007334–e1007334.

10. E X., Pickering, M.T., Debatis, M., Castillo, J., Lagadinos, A., Wang, S., Lu, S. and Kowalik, T.F. (2011) An E2F1-Mediated DNA Damage Response Contributes to the Replication of Human Cytomegalovirus. PLoS pathogens, 7, e1001342–e1001342.

11. E, X. and Kowalik, T.F. (2014) The DNA Damage Response Induced by Infection with Human Cytomegalovirus and Other Viruses. Viruses, 6, 2155–2185.

12. Luo, M.H., Rosenke, K., Czornak, K. and Fortunato, E.A. (2011) Human Cytomegalovirus Disrupts both Ataxia Telangiectasia Mutated Protein (ATM)- and ATM-Rad3-Related Kinase-Mediated DNA Damage Responses during Lytic Infection (vol 81, pg 1934, 2007). Journal of virology, 85, 3043–3043.

13. Zeltzer, S., Longmire, P., Svoboda, M., Bosco, G. and Goodrum, F. (2022) Host translesion polymerases are required for viral genome integrity. Proceedings of the National Academy of Sciences - PNAS, 119, e2203203119–e2203203119.

14. Marians, K.J. and Kornberg, R.D. (2018) Lesion Bypass and the Reactivation of Stalled Replication Forks. Annual review of biochemistry, 87, 217–238.

15. Yang, W. and Gao, Y. (2018) Translesion and Repair DNA Polymerases: Diverse Structure and Mechanism. Annual review of biochemistry, 87, 239–261.

16. Paniagua, I. and Jacobs, J.J.L. (2023) Freedom to err: The expanding cellular functions of translesion DNA polymerases. Molecular cell, 83, 3608–3621.

17. Barnes, R.P., Hile, S.E., Lee, M.Y. and Eckert, K.A. (2017) DNA polymerases eta and kappa exchange with the polymerase delta holoenzyme to complete common fragile site synthesis. DNA repair, 57, 1–11.

18. McIlwraith, M.J., Vaisman, A., Liu, Y., Fanning, E., Woodgate, R. and West, S.C. (2005) Human DNA Polymerase η Promotes DNA Synthesis from Strand Invasion Intermediates of Homologous Recombination. Molecular cell, 20, 783–792.

19. Longmire, P., Daigle, O., Zeltzer, S., Lee, M., Svoboda, M., Padilla-Rodriguez, M., Bobak, C., Bosco, G. and Goodrum, F. (2025) Complex roles for proliferating cell nuclear antigen in restricting human cytomegalovirus replication. bioRxiv, 2025.2001.2006.631530.

20. Hoege, C., Pfander, B., Moldovan, G.-L., Pyrowolakis, G. and Jentsch, S. (2002) RAD6 -dependent DNA repair is linked to modification of PCNA by ubiquitin and SUMO. Nature (London), 419, 135–141.

21. Watanabe, K., Tateishi, S., Kawasuji, M., Tsurimoto, T., Inoue, H. and Yamaizumi, M. (2004) Rad18 guides polη to replication stalling sites through physical interaction and PCNA monoubiquitination. The EMBO journal, 23, 3886–3896.

22. Huang, T.T., Nijman, S.M.B., Mirchandani, K.D., Galardy, P.J., Cohn, M.A., Haas, W., Gygi, S.P., Ploegh, H.L., Bernards, R. and D’Andrea, A.D. (2006) Regulation of monoubiquitinated PCNA by DUB autocleavage. Nat Cell Biol, 8, 341–347.

23. Cohn, M.A., Kowal, P., Yang, K., Haas, W., Huang, T.T., Gygi, S.P. and D’Andrea, A.D. (2007) A UAF1-Containing Multisubunit Protein Complex Regulates the Fanconi Anemia Pathway. Mol Cell, 28, 786–797.

24. Fox, J.T., Lee, K.-y. and Myung, K. (2011) Dynamic regulation of PCNA ubiquitylation/deubiquitylation. FEBS letters, 585, 2780–2785.

25. Moldovan, G.L. and D’Andrea, A.D. (2009) How the fanconi anemia pathway guards the genome. Annual review of genetics, 43, 223–249.

26. Ceccaldi, R., Sarangi, P. and D’Andrea, A.D. (2016) The Fanconi anaemia pathway: new players and new functions. Nature reviews. Molecular cell biology, 17, 337–349.

27. Smogorzewska, A., Matsuoka, S., Vinciguerra, P., McDonald, E.R., Hurov, K.E., Luo, J., Ballif, B.A., Gygi, S.P., Hofmann, K., D’Andrea, A.D. et al. (2007) Identification of the FANCI Protein, a Monoubiquitinated FANCD2 Paralog Required for DNA Repair. Cell, 129, 289–301.

28. Zarrella, K., Longmire, P., Zeltzer, S., Collins-McMillen, D., Hancock, M., Buehler, J., Reitsma, J.M., Terhune, S.S., Nelson, J.A. and Goodrum, F. (2023) Human cytomegalovirus UL138 interaction with USP1 activates STAT1 in infection. PLoS pathogens, 19, e1011185–e1011185.

29. Umashankar, M., Petrucelli, A., Cicchini, L., Caposio, P., Kreklywich, C.N., Rak, M., Bughio, F., Goldman, D.C., Hamlin, K.L., Nelson, J.A. et al. (2011) A Novel Human Cytomegalovirus Locus Modulates Cell Type-Specific Outcomes of Infection. PLOS PATHOG, 7, e1002444–e1002444.

30. Nguyen, C.C., Siddiquey, M.N.A., Zhang, H., Li, G. and Kamil, J.P. (2018) Human Cytomegalovirus Tropism Modulator UL148 Interacts with SEL1L, a Cellular Factor That Governs Endoplasmic Reticulum-Associated Degradation of the Viral Envelope Glycoprotein gO. Journal of virology, 92.

31. Schneider, V.A., Graves-Lindsay, T., Howe, K., Bouk, N., Chen, H.-C., Kitts, P.A., Murphy, T.D., Pruitt, K.D., Thibaud-Nissen, F., Albracht, D. et al. (2017) Evaluation of GRCh38 and de novo haploid genome assemblies demonstrates the enduring quality of the reference assembly. Genome research, 27, 849–864.

32. Langmead, B. and Salzberg, S.L. (2012) Fast gapped-read alignment with Bowtie 2. Nature methods, 9, 357–359.

33. Li, H., Handsaker, B., Wysoker, A., Fennell, T., Ruan, J., Homer, N., Marth, G., Abecasis, G. and Durbin, R. (2009) The Sequence Alignment/Map format and SAMtools. Bioinformatics, 25, 2078–2079.

34. Deatherage, D.E. and Barrick, J.E. (2014) Identification of Mutations in Laboratory-Evolved Microbes from Next-Generation Sequencing Data Using breseq. 165–188.

35. Team, R.C. (2023). R Foundation for Statistical Computing, Vienna, Austria.

36. Gu, Z., Gu, L., Eils, R., Schlesner, M. and Brors, B. (2014) circlize Implements and enhances circular visualization in R. *Bioinformatics (Oxford*, England), 30, 2811–2812.

37. Cohn, M.A., Kee, Y., Haas, W., Gygi, S.P. and D’Andrea, A.D. (2009) UAF1 Is a Subunit of Multiple Deubiquitinating Enzyme Complexes. J Biol Chem, 284, 5343–5351.

38. Li, Y., Shang, W., Xiao, G., Zhang, L.-K. and Zheng, C. (2020) A Comparative Quantitative Proteomic Analysis of HCMV-Infected Cells Highlights pUL138 as a Multifunctional Protein. *Molecules (Basel*, Switzerland), 25, 2520.

39. Nijman, S.M.B., Huang, T.T., Dirac, A.M.G., Brummelkamp, T.R., Kerkhoven, R.M., D’Andrea, A.D. and Bernards, R. (2005) The Deubiquitinating Enzyme USP1 Regulates the Fanconi Anemia Pathway. Molecular cell, 17, 331–339.

40. Garcia-Santisteban, I., Zorroza, K. and Rodriguez, J.A. (2012) Two nuclear localization signals in USP1 mediate nuclear import of the USP1/UAF1 complex. PloS one, 7, e38570.

41. Fortunato, E.A. and Spector, D.H. (1998) p53 and RPA Are Sequestered in Viral Replication Centers in the Nuclei of Cells Infected with Human Cytomegalovirus. Journal of Virology, 72, 2033–2039.

42. Penfold, M.E.T. and Mocarski, E.S. (1997) Formation of Cytomegalovirus DNA Replication Compartments Defined by Localization of Viral Proteins and DNA Synthesis. *Virology (New York*, N.Y.), 239, 46–61.

43. Strang, B.L., Boulant, S., Chang, L., Knipe, D.M., Kirchhausen, T. and Coen, D.M. (2012) Human Cytomegalovirus UL44 Concentrates at the Periphery of Replication Compartments, the Site of Viral DNA Synthesis. Journal of Virology, 86, 2089–2095.

44. Alwine, J.C. (2012) The human cytomegalovirus assembly compartment: a masterpiece of viral manipulation of cellular processes that facilitates assembly and egress. PLoS pathogens, 8, e1002878–e1002878.

45. Petrucelli, A., Rak, M., Grainger, L. and Goodrum, F. (2009) Characterization of a Novel Golgi Apparatus-Localized Latency Determinant Encoded by Human Cytomegalovirus. J Virol, 83, 5615–5629.

46. Dittmer, D. and Mocarski, E.S. (1997) Human cytomegalovirus infection inhibits G1/S transition. Journal of Virology, 71, 1629–1634.

47. Nitzsche, A., Paulus, C. and Nevels, M. (2008) Temporal Dynamics of Cytomegalovirus Chromatin Assembly in Productively Infected Human Cells. J Virol, 82, 11167–11180.

48. Lee, S.B., Lee, C.F., Ou, D.S.C., Dulal, K., Chang, L.H., Ma, C.H., Huang, C.F., Zhu, H., Lin, Y.S. and Juan, L.J. (2011) Host-viral effects of chromatin assembly factor 1 interaction with HCMV IE2. Cell research, 21, 1230–1247.

49. Manska, S. and Rossetto, C.C. (2022) Identification of cellular proteins associated with human cytomegalovirus (HCMV) DNA replication suggests novel cellular and viral interactions. *Virology (New York*, N.Y.), 566, 26–41.

50. Garcia-Higuera, I., Taniguchi, T., Ganesan, S., Meyn, M.S., Timmers, C., Hejna, J., Grompe, M. and D’Andrea, A.D. (2001) Interaction of the Fanconi Anemia Proteins and BRCA1 in a Common Pathway. Molecular cell, 7, 249–262.

51. Burssed, B., Zamariolli, M., Bellucco, F.T. and Melaragno, M.I. (2022) Mechanisms of structural chromosomal rearrangement formation. Mol Cytogenet, 15, 23.

52. Kim, H. and D’Andrea, A.D. (2012) Regulation of DNA cross-link repair by the Fanconi anemia/BRCA pathway. Genes & development, 26, 1393–1408.

53. Weitzman, M.D. and Fradet-Turcotte, A. (2018) Virus DNA Replication and the Host DNA Damage Response. Annual review of virology, 5, 141–164.

54. Kono, T., Ozawa, H. and Laimins, L. (2024) The roles of DNA damage repair and innate immune surveillance pathways in HPV pathogenesis. *Virology (New York*, N.Y.), 600, 110266.

55. Orav, M., Gagnon, D. and Archambault, J. (2019) Interaction of the human papillomavirus e1 helicase with UAF1-USP1 promotes unidirectional theta replication of viral genomes. mBio, 10, 1–15.

56. Yu, Z., Song, H., Jia, M., Zhang, J., Wang, W., Li, Q., Zhang, L. and Zhao, W. (2017) USP1-UAF1 deubiquitinase complex stabilizes TBK1 and enhances antiviral responses. The Journal of experimental medicine, 214, 3553–3563.

57. Yu, Z., Tong, L., Ma, C., Song, H., Wang, J., Chai, L., Wang, C., Wang, M., Wang, C., Yan, R. et al. (2024) The UAF1-USP1 Deubiquitinase Complex Stabilizes cGAS and Facilitates Antiviral Responses. The Journal of immunology (1950), **212**, 295-301.

58. Lahaye, X., Van, P.T., Chakraborty, C., Shmakova, A., Cao, N.T.B., Ferran, H., Ait-Mohamed, O., Maurin, M., Waterfall, J.J., Kaufer, B.B. et al. (2025) Centromeric DNA amplification triggered by viral proteins activates nuclear cGAS. Cell, 188.

59. Li, X., Stevens, P.D., Yang, H., Gulhati, P., Wang, W., Evers, B.M. and Gao, T. (2012) The deubiquitination enzyme USP46 functions as a tumor suppressor by controlling PHLPP-dependent attenuation of Akt signaling in colon cancer. ONCOGENE, 32, 471–478.

60. Gangula, N.R. and Maddika, S. (2013) WD Repeat Protein WDR48 in Complex with Deubiquitinase USP12 Suppresses Akt-dependent Cell Survival Signaling by Stabilizing PH Domain Leucine-rich Repeat Protein Phosphatase 1 (PHLPP1). J BIOL CHEM, 288, 34545–34554.

61. Buehler, J., Carpenter, E., Zeltzer, S., Igarashi, S., Rak, M., Mikell, I., Nelson, J.A. and Goodrum, F. (2019) Host signaling and EGR1 transcriptional control of human cytomegalovirus replication and latency. PLoS pathogens, 15, e1008037–e1008037.

62. Domma, A.J., Henderson, L.A., Goodrum, F.D., Moorman, N.J. and Kamil, J.P. (2023) Human cytomegalovirus attenuates AKT activity by destabilizing insulin receptor substrate proteins. Journal of virology, 97, e0056323–e0056323.

63. Mahmud, J., Geiler, B.W., Biswas, J., Miller, M.J., Myers, J.E., Matthews, S.M., Wass, A.B., O’Connor, C.M. and Chan, G.C. (2024) Delivery of US28 by incoming HCMV particles rapidly attenuates Akt activity to suppress HCMV lytic replication in monocytes. Science signaling, 17, eadn8727–eadn8727.

64. Liu, J., Jin, L., Chen, X., Yuan, Y., Zuo, Y., Miao, Y., Feng, Q., Zhang, H., Huang, F., Guo, T. et al. (2020) USP12 translocation maintains interferon antiviral efficacy by inhibiting CBP acetyltransferase activity. PLoS pathogens, 16, e1008215–e1008215.

65. Kiran, S., Dar, A., Singh, S.K., Lee, K.Y. and Dutta, A. (2018) The Deubiquitinase USP46 Is Essential for Proliferation and Tumor Growth of HPV-Transformed Cancers. Mol Cell, 72, 823–835.e825.

66. Havens, C.G. and Walter, J.C. (2011) Mechanism of CRL4, a PCNA-dependent E3 ubiquitin ligase. Genes & development, 25, 1568–1582.

67. Cukras, S., Lee, E., Palumbo, E., Benavidez, P., Moldovan, G.-L. and Kee, Y. (2016) The USP1-UAF1 complex interacts with RAD51AP1 to promote homologous recombination repair. *Cell cycle (Georgetown*, Tex.), 15, 2636–2646.

68. Williams, Samuel A., Maecker, Heather L., French, Dorothy M., Liu, J., Gregg, A., Silverstein, Leah B., Cao, Tim C., Carano, Richard A.D. and Dixit, Vishva M. (2011) USP1 Deubiquitinates ID Proteins to Preserve a Mesenchymal Stem Cell Program in Osteosarcoma. Cell, 146, 918–930.

69. Boichuk, S., Hu, L., Hein, J. and Gjoerup, O.V. (2010) Multiple DNA Damage Signaling and Repair Pathways Deregulated by Simian Virus 40 Large T Antigen. Journal of Virology, 84, 8007–8020.

70. Cherubini, G., Naim, V., Caruso, P., Burla, R., Bogliolo, M., Cundari, E., Benihoud, K., Saggio, I. and Rosselli, F. (2011) The FANC pathway is activated by adenovirus infection and promotes viral replication-dependent recombination. Nucleic acids research, 39, 5459–5473.

71. Karttunen, H., Savas, Jeffrey N., McKinney, C., Chen, Y.-H., Yates, John R., Hukkanen, V., Huang, Tony T. and Mohr, I. (2014) Co-opting the Fanconi Anemia Genomic Stability Pathway Enables Herpesvirus DNA Synthesis and Productive Growth. Molecular cell, 55, 111–122.

72. So, A., Le Guen, T., Lopez, B.S. and Guirouilh-Barbat, J. (2017) Genomic rearrangements induced by unscheduled DNA double strand breaks in somatic mammalian cells. The FEBS journal, 284, 2324–2344.

