## Supplementary Data for "Human cytomegalovirus regulates host DNA repair machinery for viral genome integrity"

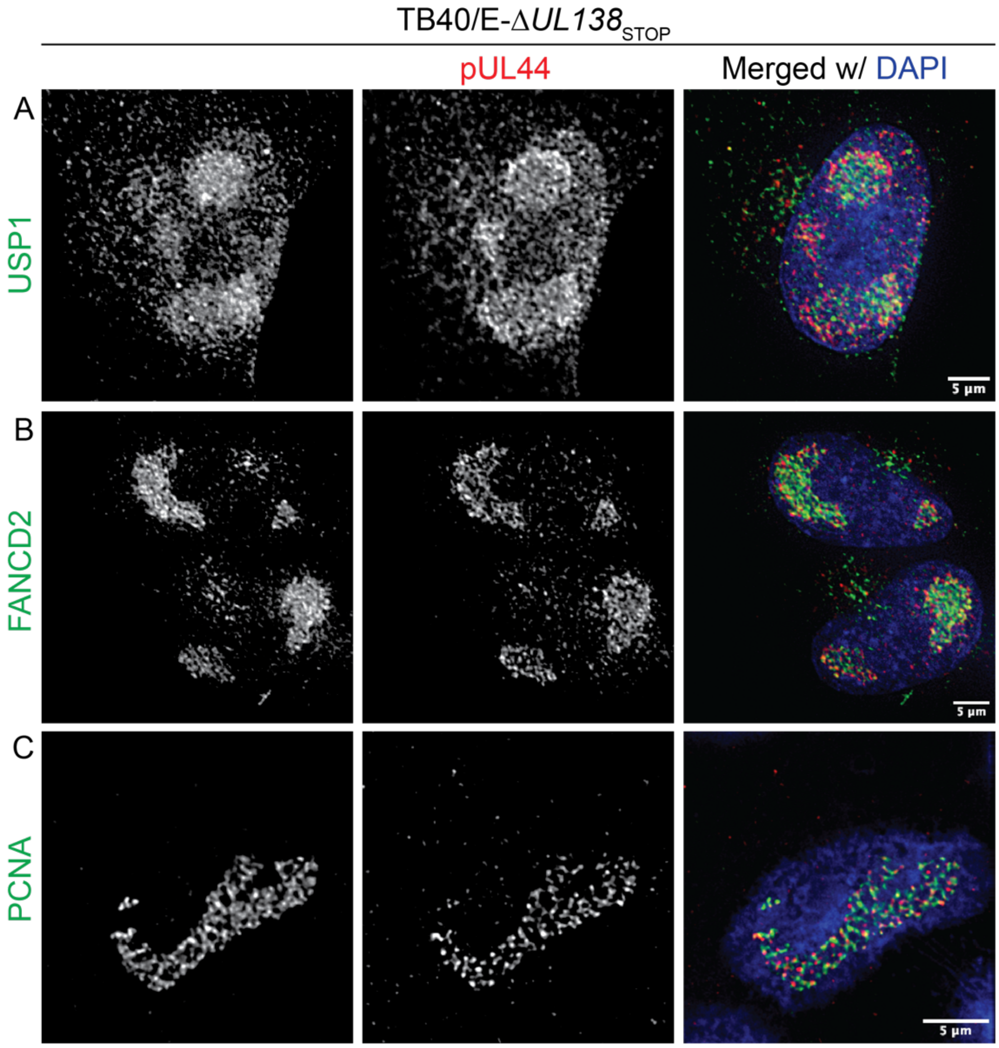


**Figure S1. USP1, FANCD2, and PCNA localization to viral RCs is independent of UL138.** Fibroblasts were infected with TB40/E-Δ*UL138*$_{STOP}$ at an MOI of 1. At 48 hpi cells were subject to CSK extraction and fixation and then processed for indirect immunofluorescence. (A) USP1, (B) FANCD2, and (C) PCNA were detected with pUL44 to mark sites of viral DNA synthesis using monoclonal antibodies specific to each and then secondary antibodies conjugated to Alexa Fluor® 546 (green) or 647 (red). Images were captured using a DeltaVision deconvolution microscope, and each deconvolved image corresponds to a single focal plane. Scale bar, 5 µm.

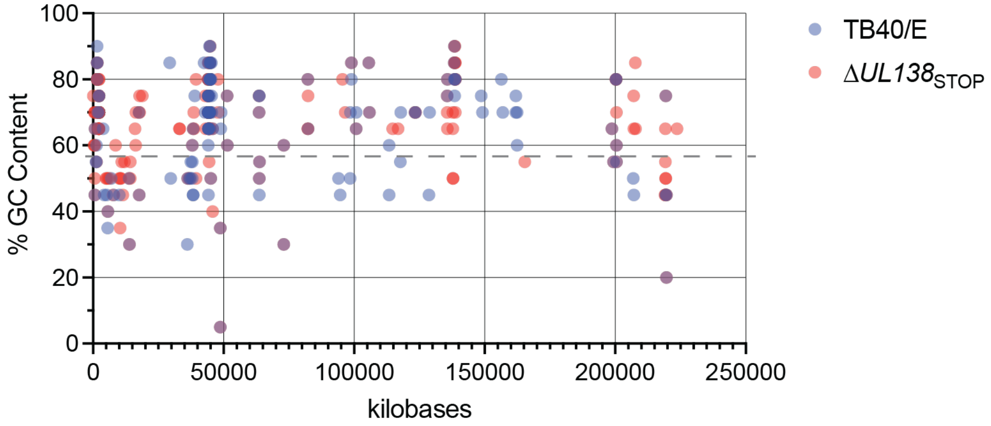


**Figure S2. GC content around novel junctions arising in TB40/E-WT and -Δ*UL138*$_{\mathbf{STOP}}$ infections.** GC content of 20-nucleotides on side of each novel junction is plotted for each infection (±SEM). Dotted line represents average 57% GC content of the whole HCMV genome.

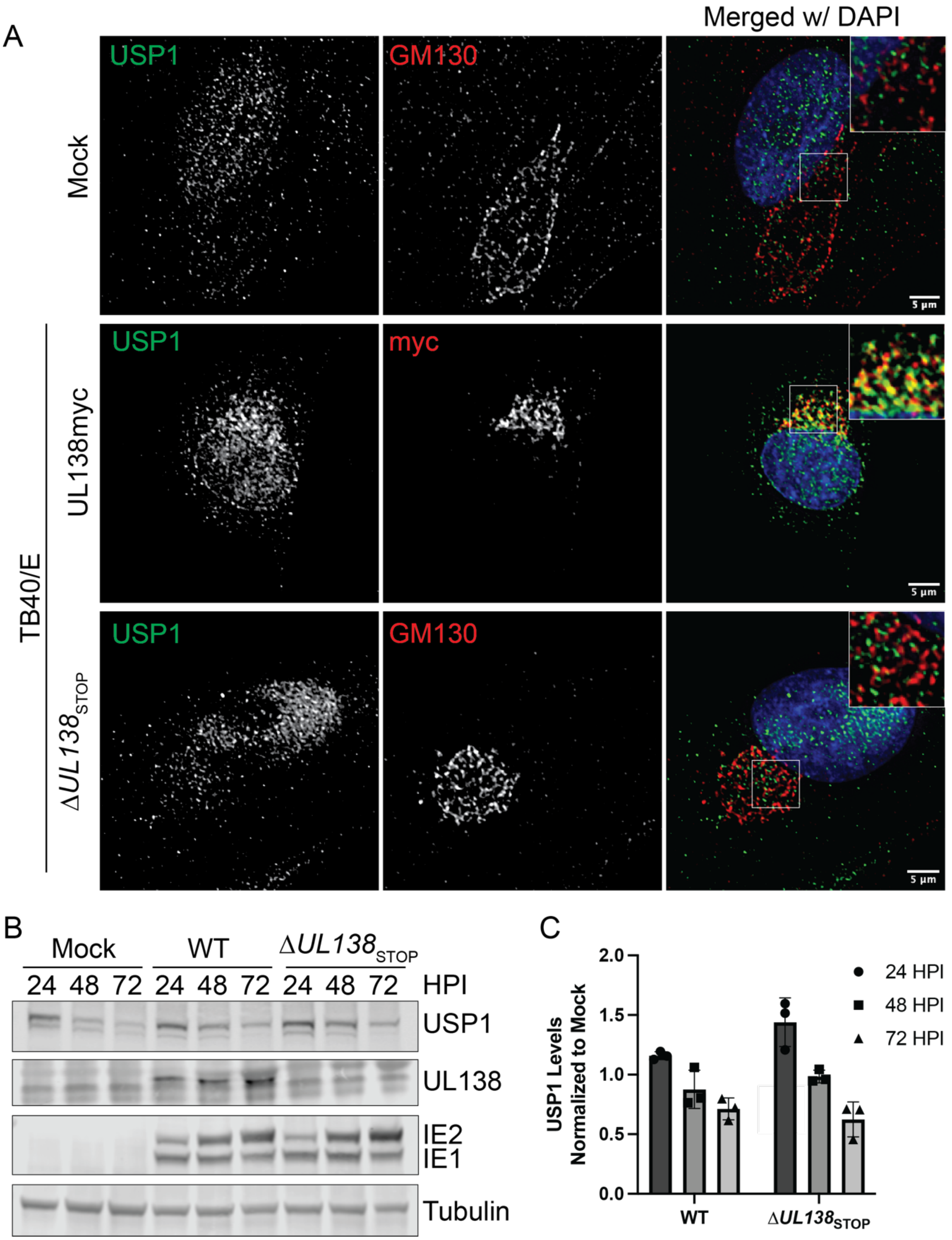
A
Merged w/ DAPI
USP1
GM130
Mock
USP1
myc
UL138myc
TB40/E
USP1
GM130
ΔUL138STOP
5 µm
B
Mock
WT
ΔUL138STOP
24 48 72 24 48 72 24 48 72
HPI
USP1
UL138
IE2
IE1
Tubulin
C
USP1 Levels Normalized to Mock
2.0
1.5
1.0
0.5
0.0
WT
ΔUL138STOP
24 HPI
48 HPI
72 HPI

**Figure S3. Investigating the effect of UL138 on USP1 localization and expression during CMV infection.**

(A) Fibroblasts were mock-infected or infected with TB40/E-UL138myc (C-terminal myc tag on UL138) or Δ*UL138*$_{STOP}$ at an MOI of 1. At 48 hpi, cells were fixed and processed for indirect immunofluorescence using monoclonal antibodies to USP1 and GM130 to detect the Golgi apparatus (where UL138 primarily localizes during infection) or myc to detect UL138. Secondary antibodies were conjugated to Alexa Fluor® 546 (green) or 647 (red). A magnified image is shown in the top right corner inset of the merged image. DAPI is used to mark the cell nucleus. Images were captured using a DeltaVision deconvolution microscope, and each deconvolved image corresponds to a single focal plane. Scale bar, 5 μm. (B) Fibroblasts were mock-infected or infected (MOI = 1) with TB40/E-WT or -Δ*UL138*$_{STOP}$ virus over a 72-hour time course. Immunoblotting was performed on whole cell lysates collected at the indicated time points. The indicated proteins were detected using antibodies specific to each and secondary antibodies conjugated to DyLight™ 680 (mouse) or 800 (rabbit). IE1/2 serve as markers for infection, and tubulin serves as a loading control. (C) Quantification of USP1 levels relative to tubulin and normalized to the Mock 24 hpi time point for WT and Δ*UL138*$_{STOP}$ infection.

| Antibody | Species | Source | Concentration |
|---|---|---|---|
| FANCD2 | Rabbit | Novus Biologicals, NB100-182 | IB: 1:10,000<br>IF: 1:200 |
| IE1/2 | Mouse | Thomas Shenk, PhD (Princeton University) | IB 1:100 |
| mUb-PCNA | Rabbit | Cell Signaling Technology (CST), #13439 | IB 1:1000<br>IF 1:100 |
| PCNA | Mouse | Santa Cruz, sc-56 | IB 1:1000 |
| PCNA | Rabbit | CST, #13110 | IF 1:400 |
| $\alpha$-Tubulin | Mouse | Sigma-Aldrich, #T9026 | IB 1:2000 |
| pUL44 | Mouse | Virusys, #CA006 | IB, IF 1:12,000 |
| UL138 | Rabbit | Open Biosystems (custom) | IB, 2.4 µg/mL |
| USP1 | Rabbit | CST, #8033 | IB: 1:1000 |
| USP1 | Rabbit | LSBio, #LS-C288484 | IF: 1:250 |
| Mouse IgG (H+L) secondary, DyLight 680 | Goat | Invitrogen, #35519 | IB: 1:6000 |
| Rabbit IgG (H+L) secondary, DyLight 800 | Goat | Invitrogen, #SA5-10036 | IB: 1:6000 |
| Rabbit IgG (H+L) secondary, Alexa Fluor 546 | Goat | Invitrogen, #A-11035 | IF 1:3000 |
| Mouse IgG (H+L) secondary, Alexa Fluor 647 | Goat | Invitrogen, #A-21236 | IF: 1:3000 |

**Table S1. Antibodies used in this study.**
